# Homeostatic Regulation of Seizure Susceptibility and Cognitive Function by Derlin-1 through Maintenance of Adult Neurogenesis

**DOI:** 10.1101/2023.04.27.538634

**Authors:** Naoya Murao, Taito Matsuda, Hisae Kadowaki, Yosuke Matsushita, Kousuke Tanimoto, Toyomasa Katagiri, Kinichi Nakashima, Hideki Nishitoh

**Author notes:** Correspondence (K.N.), (H.N.).

## Abstract

Dysfunction of organelle is closely associated with neurological diseases involving disruption of adult neurogenesis. However, the role of the endoplasmic reticulum (ER)-related molecules in this process remains largely unexplored. Here we show that Derlin-1, an ER quality controller, maintains adult neurogenesis in a spatiotemporal manner. Deletion of Derlin-1 in the mouse central nervous system induces ectopic localization of newborn neurons and impairs neural stem cells (NSCs) transition from active to quiescent states, resulting in early depletion of hippocampal NSCs. As a result, Derlin-1- deficient mice exhibit phenotypes of increased seizure susceptibility and impaired cognitive function. Reduced expression of signal transducer and activator of transcription 5b (Stat5b) was found to be responsible for the impairment of adult neurogenesis in Derlin-1-deficient NSCs. Remarkably, the inhibition of histone deacetylase activity ameliorated seizure susceptibility and cognitive dysfunction in Derlin-1-deficient mice by increasing Stat5b expression and restoring abnormal neurogenesis. Overall, our findings demonstrate that Derlin-1, as its characteristic function, plays an essential role in the homeostasis of adult neurogenesis via Stat5b expression, thus regulating seizure susceptibility and cognitive function.

---

The adult mammalian brain retains neural stem/precursor cells (NS/PCs) in restricted brain regions such as the subventricular zone (SVZ) of the lateral ventricle and the subgranular zone (SGZ) of the hippocampal dentate gyrus (DG), and these NS/PCs continuously generate neurons throughout the life of the individual (Eriksson et al. 1998; Goncalves et al. 2016). Persistent generation of neurons in the adult brain is commonly referred to as adult neurogenesis. Particularly in the DG, it plays an important role in learning and memory formation. Adult neurogenesis is disrupted in several neurological diseases associated with memory impairment (*e.g.*, seizures, depression, schizophrenia, and Alzheimer’s disease) (Snyder et al. 2011; Kang et al. 2016; Terreros-Roncal et al. 2021). Radial neural stem cells (NSCs), the source of functional neurons, are reversibly regulated to be either quiescent or proliferative (activated) by the interplay of neurogenic niche–derived signaling pathways, and this regulation of NSCs is essential for persistent neurogenesis throughout life (Bond et al. 2015; Urban et al. 2019). Furthermore, during the process of adult neurogenesis, the correct migration of newborn neurons in the adult DG is critical for physiological hippocampal function, and the mislocalization of these cells often leads to neurological dysfunction probably due to abnormal neuronal circuit formation (Scharfman and Pierce 2012; Lybrand et al. 2021). Recent studies have focused on the role of organelles such as mitochondria and lysosomes, as well as developmental signaling, transcriptional, and epigenetic pathways, in the regulation of neurogenesis (Murao et al. 2016; Beckervordersandforth et al. 2017; Kobayashi et al. 2019; Petrelli et al. 2023). However, the underlying mechanisms of the regulation of adult neurogenesis by organelles are not yet fully understood. The endoplasmic reticulum (ER) is a crucial organelle involved in the regulation of lipid and glucose metabolism, Ca^2+^ signaling, and proteostasis and has a strictly regulated quality control system in which ER-resident stress sensors recognize the accumulation of unfolded proteins and trigger the unfolded protein response (UPR). The UPR mediates the proper folding or degradation of unfolded proteins and attenuates translation to inhibit the further accumulation of proteins in the ER. Previous studies have shown that impaired ER quality contributes to the onset and exacerbation of several neurological diseases featuring learning and memory deficits (Hetz and Saxena 2017; Ghemrawi and Khair 2020). Therefore, ER function and adult neurogenesis are thought to be closely related to the mechanisms of cognitive function and neurological diseases, whereas the physiological mechanisms of these relationships remain to be elucidated.

An ER membrane protein, Derlin-1, mediates ER-associated degradation (ERAD) and ER stress–induced pre-emptive quality control (ERpQC), and is essential for ER quality control in general (Lilley and Ploegh 2004; Kadowaki et al. 2015; Kadowaki et al. 2018). We have previously shown that the interaction of Derlin-1 with amyotrophic lateral sclerosis (ALS)-associated superoxide dismutase 1 (SOD1) mutants leads to a pathological UPR and motor neuron dysfunction (Nishitoh et al. 2008). Furthermore, loss of Derlin-1 in the central nervous system (CNS) induces brain atrophy and motor dysfunction by impairing neuronal cholesterol biosynthesis, which is regulated on the ER membrane (Sugiyama et al. 2021). Chemical chaperones such as 4-phenylbutyric acid (4-PBA) can rescue the above phenotypes in Derlin-1-deficient mice, indicating that Derlin-1-mediated ER quality control is essential for brain development and function (Sugiyama et al. 2022). Derlin-1 is expressed in adult hippocampal NSCs, and its expression fluctuates across the stages of adult NSCs (Shin et al. 2015). Considering these findings, we hypothesize that Derlin-1 in NSCs may be necessary for the regulation of adult neurogenesis and related behaviors.

Here, we show that Derlin-1 is responsible for the regulation of adult neurogenesis in a spatiotemporal manner, i.e., the transition of NSCs from active to quiescent states and the localization and survival of newborn neurons, and that NSCs are depleted early in mice with CNS-specific Derlin- 1 deficiency. Furthermore, the loss of *Derlin-1* (*Derl1*) in mice increases seizure susceptibility and impairs cognitive function. Signal transducer and activator of transcription 5b (Stat5b) was identified as a regulator of adult neurogenesis downstream of Derlin-1. Surprisingly, 4-PBA rescues the phenotype of Derlin-1-deficient mice via its inhibitory action on histone deacetylase (HDAC), not its chaperone activity. Overall, our work demonstrates that the Derlin-1-Stat5b axis is essential for the homeostasis of adult neurogenesis and consequently plays an important role in regulating seizure susceptibility and cognitive function.

## Results

### Abnormal proliferation of NSCs due to Derlin-1 deficiency

Impaired ER function contributes to the pathogenesis of several neurological diseases characterized by cognitive dysfunction, and Derlin-1 expression in the mouse hippocampus is varies across the stages of adult NSCs (Shin et al. 2015; Hetz and Saxena 2017; Ghemrawi and Khair 2020). To understand the role of Derlin-1 in adult neurogenesis, we analyzed mice with CNS-specific Derlin-1 deficiency (*Derl1^NesCre^* mice) generated by mating *Derl1^flox/flox^* (*Derl1^f/f^*) mice harboring a floxed *Derl1* gene with transgenic mice expressing Cre recombinase under the control of the *nestin* promoter [*Tg(Nes-Cre)1Kag* mice] (Isaka et al. 1999; Sugiyama et al. 2021). Derlin-1 protein was barely detectable in the hippocampal region and DG of *Derl1^NesCre^* mice (Supplemental Fig. S1A). In the DG of *Derl1^NesCre^* mice, we confirmed the induction of ER stress using DNA microarrays and gene set enrichment analysis (GSEA) focused on the term “Response to ER stress” (Supplemental Fig. S1B), consistent with our previous findings that ER stress is induced in the cerebellum of this mouse line (Sugiyama et al. 2021). In the mouse DG and granular cell layer (GCL), structures are fully formed at approximately two weeks of age, and the number of neural progenitor cells in the molecular layer decreases as DG development proceeds (Noguchi et al. 2016). We examined whether Derlin-1 deficiency affects hippocampal development by observing the GCL morphology and the number and localization of Tbr2- and Ki67-positive cells; we found no change in the DG during development (Supplemental Fig. S1C ‒ G). Next, we injected 8-week-old *Derl1^f/f^*or *Derl1^NesCre^* mice with bromodeoxyuridine (BrdU) once a day for 7 days to examine the effects of Derlin-1 deficiency on adult hippocampal neurogenesis (Fig. 1A). The numbers of BrdU-positive proliferating cells and BrdU- and DCX-double-positive newborn neurons were increased in *Derl1^NesCre^* mice (Fig. 1B‒D), suggesting that Derlin-1-deficient NSCs excessively proliferate within a time frame of one week in the adult hippocampus.

**Figure 1.**
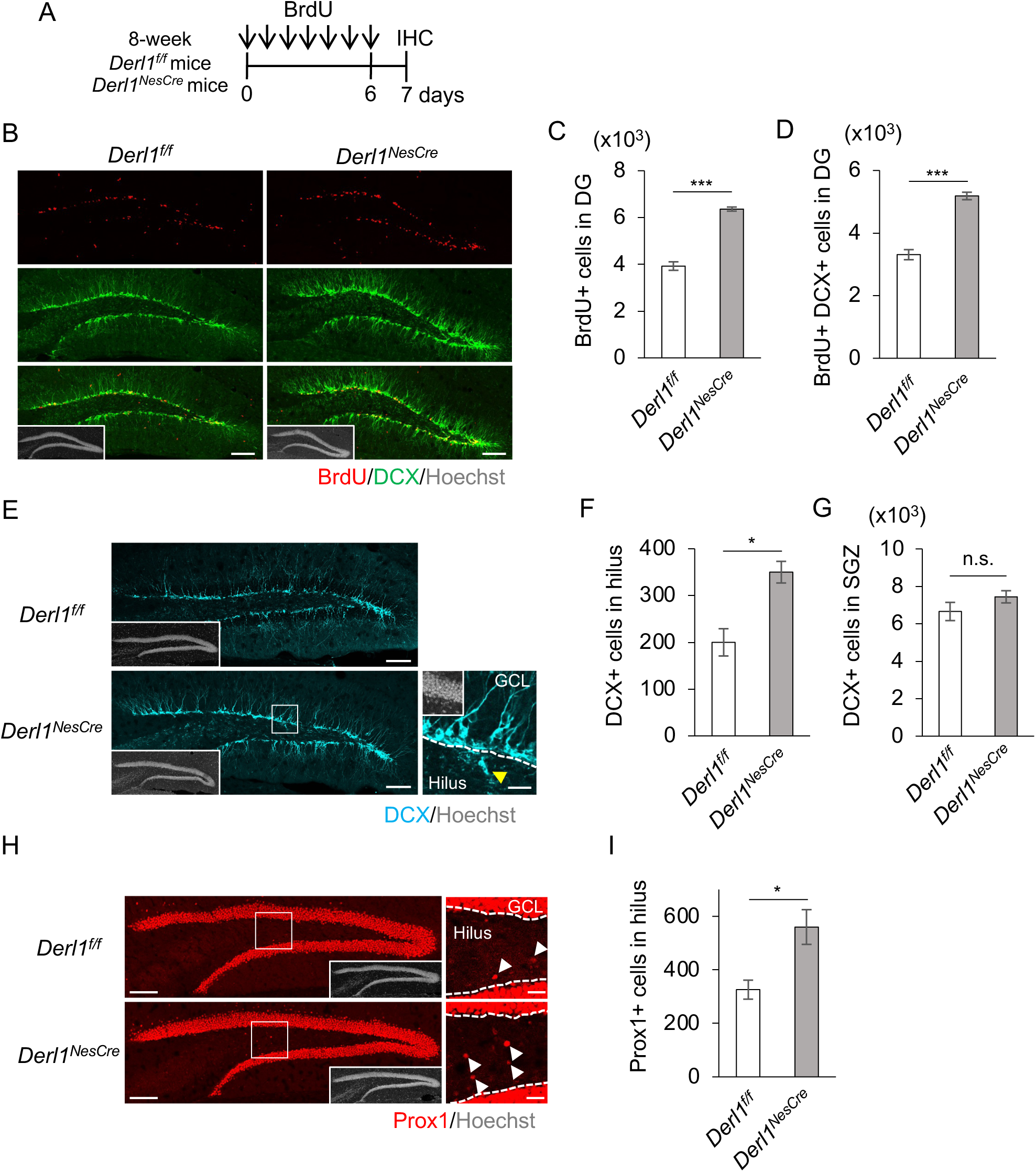
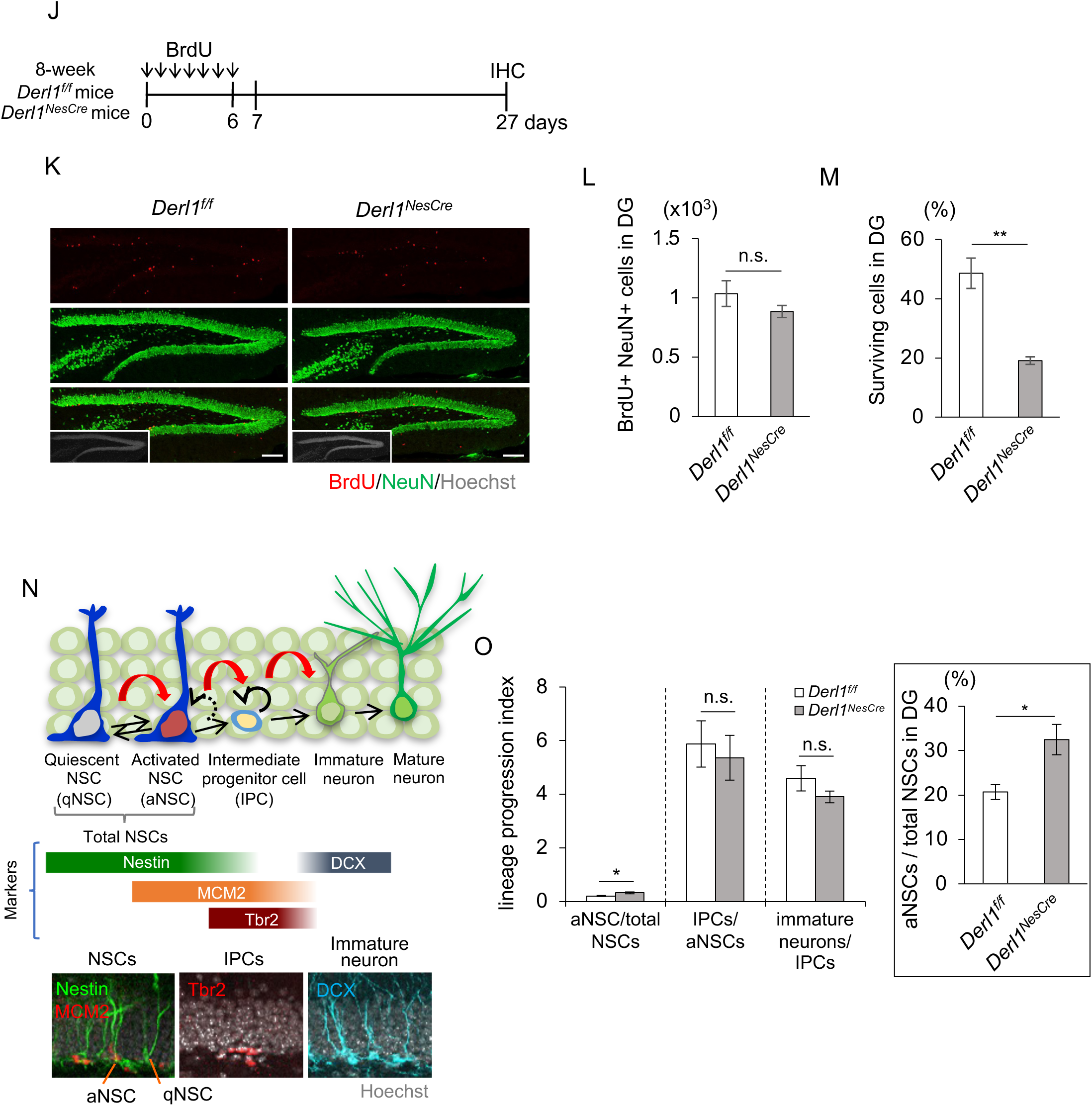
Loss of *Derl1* perturbs adult hippocampal neurogenesis. (A) Experimental scheme for investigating the proliferation of NS/PCs and neurogenesis in *Derl1^f/f^* and *Derl1^NesCre^* mice. (B) Representative immunofluorescence images of DG staining for BrdU (red), DCX (green), and Hoechst (gray; insets). Scale bars: 100 μm. (*C*,*D*) Quantification of the numbers of BrdU-positive (BrdU+) proliferating cells (*C*) and BrdU+ DCX+ newborn immature neurons (*D*) in the DG of *Derl1^f/f^* and *Derl1^NesCre^* mice (n = 3 mice). (*E*) Representative immunofluorescence images of DG staining for DCX (cyan) and Hoechst (gray; insets). The areas outlined by a white rectangle of the *Derl1^NesCre^* panel are enlarged to the right. The yellow arrowhead indicates an ectopic DCX+ immature neuron in the hilus, and dashed white lines indicate the boundaries between the GCL and the hilus. Scale bars, 100 μm (left) and 20 μm (right). (*F*,*G*) Quantification of the number of DCX+ cells in the hilus (*F*) and SGZ (*G*) of *Derl1^f/f^* and *Derl1^NesCre^* mice (n = 3 mice). (H) Representative immunofluorescence images of the DG with Prox1 (red) and Hoechst staining (gray; insets). The areas outlined by a white rectangle are enlarged to the right. The white arrowheads indicate Prox1+ ectopic neurons in the hilus, and the dashed white lines indicate the boundaries between the GCL and the hilus. Scale bars, 100 μm (left) and 20 μm (right). (I) Quantification of the number of Prox1+ cells in the hilus of *Derl1^f/f^* and *Derl1^NesCre^* mice (n = 3 mice). (J) Experimental scheme for assessing neurogenesis in the DG of *Derl1^f/f^* and *Derl1^NesCre^* mice. (K) Representative immunofluorescence images of the DG with BrdU (red), NeuN (green), and Hoechst staining (gray; insets). Scale bars: 100 μm. (L) Quantification of the number of BrdU+ NeuN+ newborn mature neurons in the DG of *Derl1^f/f^* and *Derl1^NesCre^* mice (n = 6; *Derl1^f/f^* mice, n = 8; *Derl1^NesCre^* mice). (M) The percentage of BrdU+ cells surviving between 1 and 3 weeks in the DG of *Derl1^f/f^* and *Derl1^NesCre^* mice. The survival ratio was obtained by dividing the total number of BrdU+ cells at 3 weeks (day 27 overall) by the total number at 1 day (day 7 overall) after the last BrdU injection (n = 3 mice). (N) Schematic diagram of the developmental stage of adult neurogenesis (top panel) and specific marker proteins for each stage (middle panel). Representative immunofluorescence images of Nestin+ MCM2- (qNSC), Nestin+ MCM2+ (aNSC), Tbr2+ (IPC), and DCX+ (immature neuron) staining (bottom panel). (O) Comparison of the lineage progression index between *Derl1^f/f^*and *Derl1^NesCre^* mice. The lineage progression index is calculated by dividing the number of cells at a defined developmental stage by the number of cells at the preceding developmental stage (aNSCs normalized to total NSCs, IPCs normalized to aNSCs, immature neurons normalized to IPCs) (n = 3 mice). The inset shows the proportion of the number of aNSCs to total NSCs (n = 3 mice). Bar graphs are presented as the mean ± SEM. *P < 0.05, **P < 0.01, and ***P < 0.001 by Student’s t test. n.s., not significant.

The Derlin family consists of Derlin-1, 2, and 3. Derlin-1 and 2 are ubiquitously expressed, including in the brain, while Derlin-3 is not, and both Derlin-1 and Derlin-2 play important roles in brain development and function by maintaining ER quality (Nishitoh et al. 2008; Dougan et al. 2011; Sugiyama et al. 2022). We generated *Derl2^NesCre^*mice and injected BrdU once a day for 7 days in 8- week-old *Derl2^NesCre^* and control (*Derl2^f/f^*) mice (Supplemental Fig. S1H). The expression of ER stress–responsive genes was significantly increased in the DG of *Derl2^NesCre^* mice (Fig. 1I), but surprisingly, there was no significant difference in the number of BrdU-positive cells or BrdU- and DCX-double-positive cells (Supplemental Fig. S1J‒L). These results suggest that the disturbance of adult neurogenesis in *Derl1^NesCre^*mice is independent of ER stress. We further examined whether Derlin-1 expression in neurons is involved in adult neurogenesis. *Derl1^CaMKIIαCre^* mice, in which *Derl1* is specifically deleted in neurons, showed no changes in cell proliferation and DCX-positive neuron production (Supplemental Fig. S1H,M‒O). We next focused on the number and location of newborn neurons in *Derl1^NesCre^* mice, as the number and localization of newborn neurons are important in the developmental process of adult neurogenesis. In the DG of *Derl1^NesCre^* mice, the numbers of both DCX-positive immature neurons (Fig. 1E,F) and mature neurons positive for the granular cell marker Prox1 (Fig. 1H,I) located in the hilus were increased. In contrast, the number of DCX-positive cells in the SGZ was not changed (Fig. 1G). Collectively, loss of *Derl1* induces ectopic neurogenesis. To examine whether the abnormally generated *Derl1^NesCre^*neurons mature, *Derl1^NesCre^* and control mice were analyzed 3 weeks after 7 days of BrdU injection (Fig. 1J). BrdU- and NeuN-double-positive mature neurons were not increased (Fig. 1K,L) and the survival ratio of newborn neurons was significantly decreased in the DG of *Derl1^NesCre^*mice (Fig. 1M), suggesting that Derlin-1-deficient NSCs differentiate into immature neurons but do not become mature neurons. To identify the abnormality in the stage of adult neurogenesis in *Derl1^NesCre^* mice, we investigated the lineage progression index of each developmental stage using cellular markers of specific developmental stages; we calculated this index by dividing the number of cells of a defined developmental stage by the number of cells of the preceding developmental stage (Fig. 1N,O). Intriguingly, we discovered that the lineage progression index of activated NSCs, but not intermediate progenitor cells or immature neurons, was specifically increased during the developmental stage of adult neurogenesis in *Derl1^NesCre^* mice (Fig. 1O). The percentage of activated NSCs among total NSCs was increased by more than 10% in *Derl1^NesCre^* mice (Fig. 1O, inset). These results suggest that Derlin-1 primarily regulates the quiescent and active states of NSCs and is also involved in the localization and survival of newborn neurons.

### Enhanced seizure susceptibility due to Derlin-1 deficiency

Ectopic localization of neurons in the hilus of the DG is frequently observed in patients with temporal lobe epilepsy, the most common form of epilepsy in adults, and in animal models of this disease (Parent et al. 2006; Hester and Danzer 2013; Cho et al. 2015; Matsuda et al. 2015). These ectopic neurons are more excitable than neurons in the GCL (Zhan et al. 2010; Cameron et al. 2011). Having observed that the number of Prox1-positive neurons in the hilus is increased in *Derl1^NesCre^* mice compared to control mice, we investigated the seizure susceptibility of *Derl1^NesCre^*mice. Kainic acid (KA), an agonist for a subtype of ionotropic glutamate receptor, was administered to 2-month-old *Derl1^NesCre^* and *Derl1^f/f^* control mice, and the seizure phenotype was observed for 1 h. *Derl1^NesCre^* mice showed higher seizure scores than control mice (Fig. 2A,B). These data suggest that Derlin-1 contributes to the appropriate localization of newly generated neurons, which is important in reducing seizure susceptibility.

**Figure 2.**
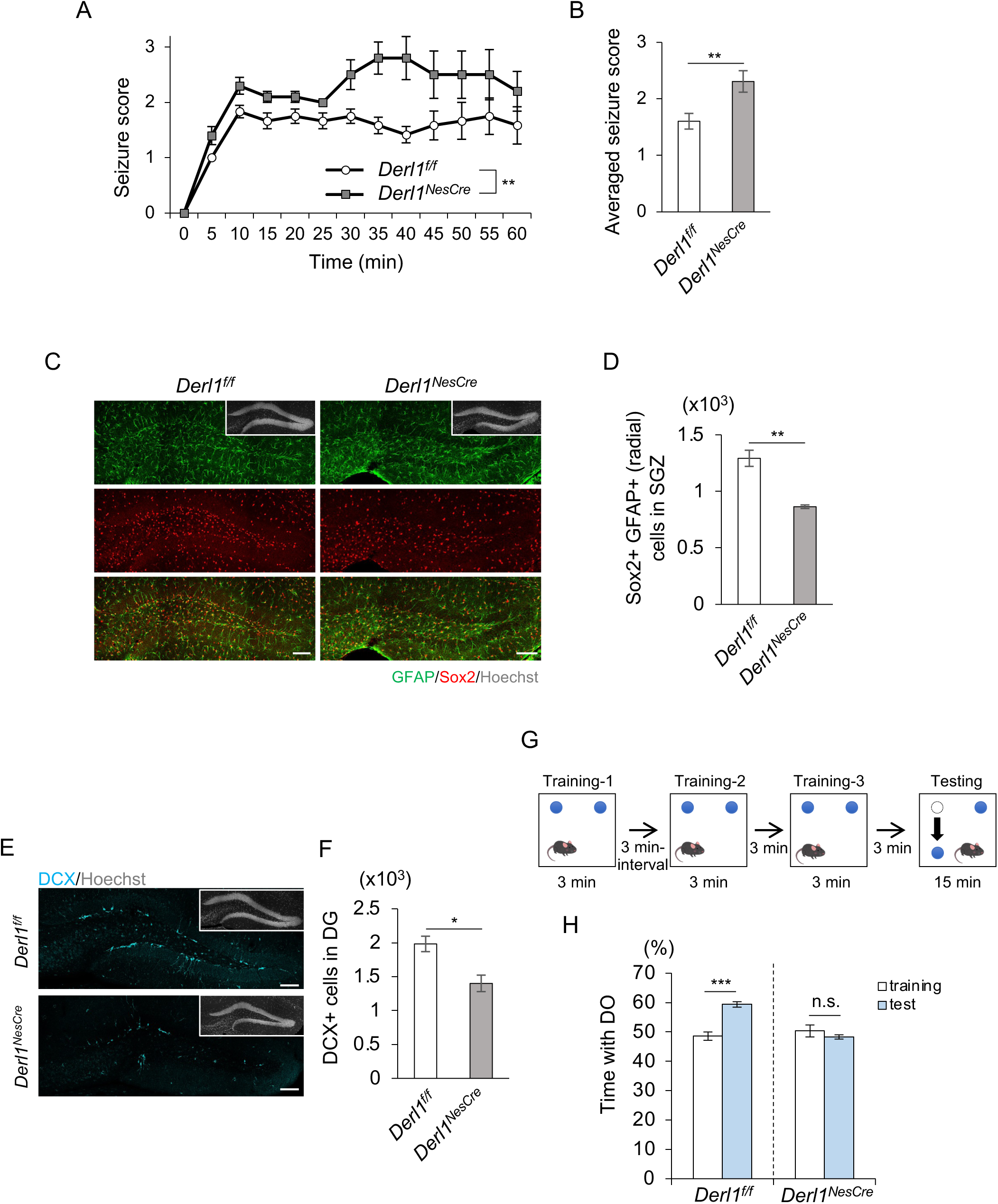
Derlin-1 deficiency increases seizure susceptibility and impairs cognitive function. (A) Time plot showing the mean seizure score over 1 h after KA treatment in *Derl1^f/f^* and *Derl1^NesCre^* mice (n = 12; *Derl1^f/f^* mice, n = 10; *Derl1^NesCre^* mice). (B) Bar graph showing averaged seizure scores in *Derl1^f/f^* and *Derl1^NesCre^* mice. (C) Representative immunofluorescence images of the DG with GFAP (green), Sox2 (red), and Hoechst staining (gray; insets). Scale bars: 100 μm. (D) Quantification of the number of Sox2+ radial GFAP+ NSCs in the SGZ of 9-month-old *Derl1^f/f^* and *Derl1^NesCre^* mice (n = 3 mice). (E) Representative immunofluorescence images of the DG stained for DCX (cyan) in 9-month-old *Derl1^f/f^* and *Derl1^NesCre^* mice. Scale bars: 100 μm. (F) Quantification of the number of DCX+ immature neurons in the DG of 9-month-old *Derl1^f/f^* and *Derl1^NesCre^* mice (n = 4 mice). (G) Schematic diagram of the experimental protocol for the novel location recognition test. (H) Percentage of time spent with the displaced object (DO) during the training and testing phases in 4-month-old *Derl1^f/f^* and *Derl1^NesCre^* mice (n = 10; *Derl1^f/f^* mice, n = 6; *Derl1^NesCre^* mice). Bar graphs are presented as the mean ± SEM. *P < 0.05, **P < 0.01, and ***P < 0.001 by two-way repeated-measures ANOVA (*A*) or Student’s t test (*B*, *D*, *F* and *H*). n.s., not significant.

### Requirement of Derlin-1 for maintenance of NSC numbers and cognitive function in the aged mouse brain

To examine whether the increase in active NSCs in *Derl1^NesCre^*mice affects the maintenance of neurogenesis throughout life, we quantified the number of NSCs in middle-aged (9-month-old) mice. In *Derl1^NesCre^* mice, the number of NSCs in the DG was markedly decreased compared to that in control mice (Fig. 2C,D). The number of DCX-positive immature neurons was also decreased in *Derl1^NesCre^*mice (Fig. 2E,F). On the basis of these results together, it is conceivable that Derlin-1 is required to maintain the NSC pool in the aged mouse brain and to ensure adequate neurogenesis successively throughout life. Adult hippocampal neurogenesis is vital for cognitive function, and the novel location recognition test, using spatial discrimination ability as an index, is known to reflect hippocampus-dependent cognitive function (Goodman et al. 2010; Goncalves et al. 2016). Four-month-old *Derl1^NesCre^* mice showed a reduced preference for the displaced object (DO) in the testing phase (Fig. 2G,H), suggesting that they were unable to identify changes in the locations of objects. In contrast, *Derl2^NesCre^* mice with unchanged adult neurogenesis spent more time with the DO in the testing phase, similar to control mice (Supplemental Fig. S2). These results suggest that hippocampus-dependent cognitive function is impaired in *Derl1^NesCre^* mice due to disrupted adult neurogenesis.

### Requirement of Derlin-1 for the transition of NSCs from active to quiescent states

To elucidate the mechanism by which the ratio of activated NSCs increases in the DG of *Derl1^NesCre^* mice (Fig. 1O), we employed cultured adult rat hippocampal NSCs. Adult rat hippocampal NSCs have been reported to remain in a highly proliferative state when treated with basic fibroblast growth factor (bFGF), and treatment with diazepam or BMP4 artificially induces NSCs into a quiescent state (Mira et al. 2010; Mukherjee et al. 2016; Doi et al. 2021). We used these culture conditions to examine the effect of Derlin-1 deficiency on the transition of NSCs from active to quiescent states. Adult rat hippocampal NSCs transfected with anti-Derl1 siRNA were cultured for 2 days in proliferation medium, then for another 2 days in proliferation medium or diazepam- or BMP4-containing quiescence medium, and analyzed after 30 min of 5-ethynyl-2-deoxyuridine (EdU) treatment (Fig. 3A). The percentage of EdU-positive proliferating NSCs in *Derl1* knockdown (siDerl1) NSCs was unchanged under proliferating conditions compared to control (siControl) NSCs (Fig. 3C). By contrast, the percentage of EdU-positive proliferating siDerl1 NSCs was increased in quiescent conditions (Fig. 3B,C). In the adult hippocampal DG, activated NSCs are known to return to a quiescent state at a certain rate (Harris et al. 2021). Therefore, our results from *in vitro* experiments suggest that the increased percentage of activated NSCs among total NSCs in *Derl1^NesCre^*mice may be due to a defect in the Derlin-1-mediated transition of NSCs from active to quiescent states. To investigate the possibility that secreted factors from Derlin-1-deficient NSCs inhibit the transition from active to quiescent states, the culture medium of activated wild-type NSCs was replaced with 50% volume of culture medium from siControl NSCs or siDerl1 NSCs and 50% volume of new quiescence medium (Supplemental Fig. S3A). The proliferating cell ratio was unchanged (Supplemental Fig. S3B,C), suggesting that Derlin-1 deficiency would not result in the secretion of factors that dominantly inhibit the transition from active to quiescent states. We next examined whether factors secreted by wild-type NSCs during quiescent state induction were sufficient to improve inhibition of the transition from active to quiescent states in Derlin-1-deficient NSCs. (Supplemental Fig. S3C). The percentage of proliferating NSCs in siDerl1 NSCs remained higher than that in siControl NSCs, even when 50% volume of culture medium from wild-type NSCs was used (Supplemental Fig. S3D). These data suggest that Derlin-1 regulates the transition of NSCs from active to quiescent states primarily through a cell-autonomous manner.

**Figure 3.**
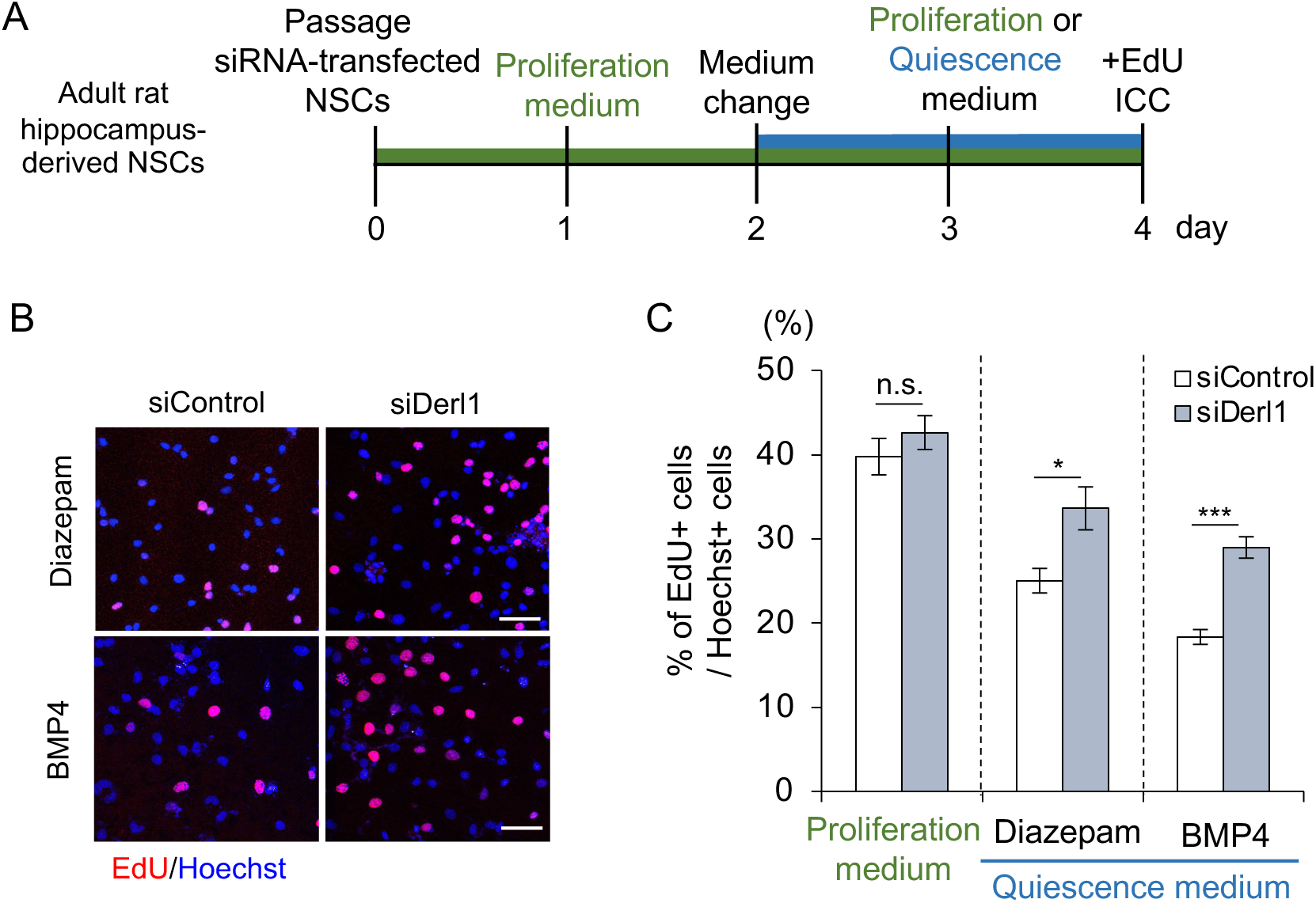
Derlin-1 is required for the transition of NSCs from active to quiescent states. (A) Experimental scheme to induce the transition of control and *Derl1* knockdown NSCs from active to quiescent states. (B) Representative images of EdU (red) and Hoechst (blue) staining in control (siControl) and *Derl1* knockdown (siDerl1) NSCs after induction of quiescence with diazepam or BMP4 for 2 days. NSCs were fixed 30 min after the addition of EdU. Scale bars: 50 μm. (C) Quantification of the percentage of EdU+ proliferating NSCs among total Hoechst+ cells in siControl and siDerl1 NSCs with proliferative conditions or induction of quiescence with diazepam or BMP4 for 2 days (n = 6; proliferation medium and diazepam condition, n = 8; BMP4 condition). Bar graphs are presented as the mean ± SEM. *P < 0.05 and ***P < 0.001 by Student’s t test. n.s., not significant.

### Requirement of Stat5b expression for maintenance of NSCs

To understand the mechanism by which Derlin-1 deficiency impairs the transition of NSCs from active to quiescent states, RNA sequencing (RNA-seq) was performed on adult rat hippocampal NSCs induced to enter the quiescent state (Supplemental Fig. S4A). In siDerl1 NSCs, the expression levels of 184 genes were significantly increased (>1.5-fold), and those of 180 genes were significantly decreased (<0.8-fold) compared to siControl NSCs (Supplemental Table S1). Although Derlin-1 deficiency increased the expression of ER stress-related genes in the DG (Supplemental Fig. S1B), significant enrichment of ER stress-related genes among downstream targets of Derlin-1 was not observed in adult rat hippocampal NSCs (Supplemental Fig. S4B). Therefore, it is conceivable that the abnormal transition of siDerl1 NSCs from active to quiescent states may not be triggered by ER stress itself but rather by changes in unconventional genes regulated by Derlin-1. Since it is well known that many transcription factors regulate the expression of genes involved in stem cell states, the group of genes whose expression is altered by Derlin-1 deficiency was searched in the bracket of transcription factors (Supplemental Table S1) (Sarkar and Hochedlinger 2013; Takashima and Suzuki 2013). The expression levels of 6 transcription factors were increased in siDerl1 NSCs, while those of 9 transcription factors were decreased (Fig. 4C). Among these transcription factors, Stat5b has been reported to be involved in the maintenance of tissue stem cell quiescence (Wang et al. 2009; Wang et al. 2019; Kollmann et al. 2021). The expression of *Stat5b* was decreased in siDerl1 NSCs in both the proliferative and quiescent states (Fig. 4A,B). Additionally, the expression of Stat5b protein was confirmed to be lower in siDerl1 NSCs than in siControl NSCs (Fig. 4C). These findings suggest that the expression of Stat5b is transcriptionally regulated downstream of Derlin-1. We hypothesized that decreased expression of Stat5b may be responsible for the disruption of homeostasis in siDerl1 NSCs. To test this hypothesis, adult rat hippocampal NSCs transfected with an siRNA against *Stat5b* were cultured for 2 days in proliferation medium and for 2 days in quiescence medium containing BMP4 (Fig. 4A). Intriguingly, we found that Stat5b deficiency increased the percentage of EdU- or Ki67- positive proliferating adult rat hippocampal NSCs (Fig. 4D‒F). This increase in the percentage of EdU-positive proliferating cells after induction of quiescence was consistent with the results from siDerl1 NSCs (Fig. 3B,C). To examine whether the expression of Stat5b is sufficient to restore the impaired transition of siDerl1 NSCs, NSCs infected with a control lentivirus encoding Venus (a GFP variant) or a lentivirus encoding Venus-tagged Stat5b were induced to enter the quiescent state (Fig. 4G). We found that exogenously expressed Stat5b restored the percentage of proliferating siDerl1 NSCs to that of proliferating siControl NSCs (Fig. 4H,I), suggesting that Stat5b expressed downstream of Derlin-1 regulates the transition of NSCs from active to quiescent states.

**Figure 4.**
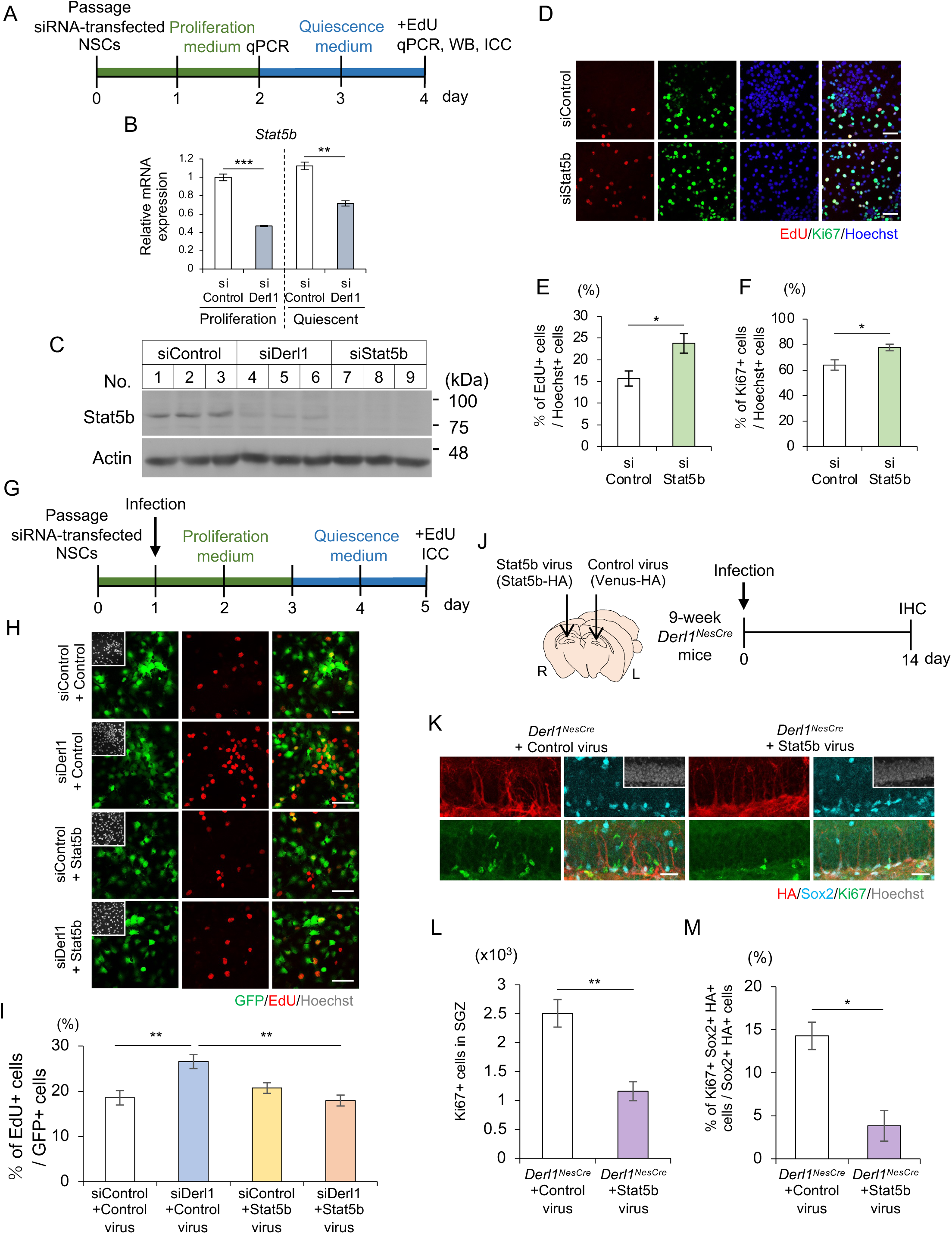
Stat5b is required for the transition of NSCs from active to quiescent states. (A) Experimental scheme for evaluating the relevance of Stat5b underlying the impairment of NSC transition to quiescence by *Derl1* knockdown. (B) Expression of *Stat5b* in siControl and siDerl1 NSCs under proliferation and quiescent conditions. Gene expression levels were estimated by qPCR and normalized to that of *β-actin* (n = 3). (C) Representative immunoblots (IB) of siControl, siDerl1 and siStat5b NSCs after induction of quiescence. Whole-cell lysates were analyzed by IB with Stat5b and actin antibodies. (D) Representative images of EdU (red), Ki67 (green), and Hoechst (blue) staining in siControl and siStat5b NSCs after induction of quiescence with BMP4 for 2 days. NSCs were fixed 30 min after the addition of EdU. Scale bars: 50 μm. (*E*,*F*) Quantification of the percentages of EdU+ (*E*) and Ki67+ (*F*) proliferating NSCs among total Hoechst+ cells in siControl and siStat5b NSCs after induction of quiescence with BMP4 for 2 days (n = 3). (G) Experimental scheme for investigating the requirement of Stat5b for impairment of NSC transition to quiescence by *Derl1* knockdown. (H) Representative images of GFP (green), EdU (red), and Hoechst (gray; insets) staining in siControl and siDerl1 quiescence-conditioned NSCs with or without exogenous Stat5b expression. NSCs were fixed 30 min after the addition of EdU. Scale bars: 50 μm. (I) Quantification of the percentage of EdU+ proliferating NSCs among total GFP+ cells in siControl and siDerl1 NSCs with or without exogenous Stat5b expression (n = 4). (J) Experimental scheme for assessing the effect of Stat5b expression in the DG on NSC proliferation in *Derl1^NesCre^* mice. (K) Representative immunofluorescence images of the DG with HA (red), Sox2 (cyan), Ki67 (green), and Hoechst staining (gray; insets). Scale bars: 25 μm. (L) Quantification of the number of Ki67+ proliferating cells in the DG of *Derl1^NesCre^* mice with or without exogenous Stat5b expression (n = 3 mice). (M) Quantification of the percentage of Ki67+ Sox2+ HA+ proliferating NS/PCs among total Sox2+ HA+ cells in the DG of *Derl1^NesCre^* mice with or without exogenous Stat5b expression (n = 3 mice). Bar graphs are presented as the mean ± SEM. *P < 0.05, **P < 0.01, and ***P < 0.001 by Student’s t test (*B*, *E*, *F*, *L* and *M*) or one-way ANOVA followed by Tukey’s test (*I*).

Stat5b is a member of the Stat family of proteins, which are phosphorylated by receptor-bound Janus kinase (JAK) in response to cytokines and growth factors to form homodimers or heterodimers that translocate into the nucleus to act as transcriptional activators (Able et al. 2017). To examine whether the transcriptional activity of Stat5b is required for the restoration of the impaired transition of siDerl1 NSCs from active to quiescent states, NSCs were infected with a virus encoding mutant Stat5b (Y699F), in which the tyrosine phosphorylation sites required for activation were replaced with phenylalanine. We found that exogenously expressed Stat5b (Y699F) also reduced the abnormal proliferation of siDerl1 NSCs (Supplemental Fig. S4D). Moreover, downstream target genes of Stat5b were not enriched among the downstream targets of Derlin-1 in adult rat hippocampal NSCs (Fig. 4E), suggesting that the transcriptional activity of Stat5b may not be required for the maintenance of NSCs. We next examined whether exogenous expression of Stat5b suppresses the abnormal proliferation of Derlin-1-deficient NSCs *in vivo*. The DG of the hippocampus in 9-week-old *Derl1^NesCre^* mouse brains was infected by intracerebral injection with a lentivirus encoding control Venus on one side and a lentivirus encoding Stat5b on the other side, and the mice were analyzed by immunostaining after 2 weeks (Fig. 4J). Compared with the control virus–infected DG, the DG infected with the Stat5b-encoding virus had a reduced number of Ki67-positive proliferating cells, as well as a reduced percentage of proliferating NS/PCs among virus–infected HA-positive NS/PCs (Fig. 4K‒M). On the basis of these results together, it is suggested that the reduced expression of Stat5b is responsible for the abnormal proliferation of NSCs in *Derl1^NesCre^* mice.

### Restoration of impaired transition of Derlin-1-deficient NSCs from active to quiescent states by 4-PBA

We have previously reported that continuous treatment of *Derl1^NesCre^* mice with 4-PBA improved motor impairment due to brain atrophy (Sugiyama et al. 2021). 4-PBA acts not only as a chemical chaperone but also as an HDAC inhibitor, and other HDAC inhibitors, such as valproic acid (VPA), counteract neurological diseases including spinal cord injury and hearing loss in mouse models by promoting neuronal differentiation (Abematsu et al. 2010; Kusaczuk et al. 2015; Wakizono et al. 2021). Based on these findings, to verify the possibility that 4-PBA may be effective in rescuing the abnormality of Derlin-1-deficient NSCs, NSCs were pretreated with 4-PBA one day before induction to a quiescence state, and the percentage of proliferating NSCs was quantified 3 days later (Fig. 5A). The percentage of proliferating siDerl1 NSCs was significantly reduced by 4-PBA treatment (Fig. 5B,C). We examined the effect of 4-PBA treatment on *Stat5b* expression in NSCs (Fig. 5D). The expression of *Stat5b* was increased in siControl and siDerl1 NSCs by 4-PBA treatment, and the reduced *Stat5b* expression in siDerl1 NSCs recovered to the same level found in the vehicle-treated siControl NSCs (Fig. 5E). We next examined whether this induction of *Stat5b* expression depends on chaperone activity or HDAC inhibition activity by using other chemical chaperones, tauroursodeoxycholic acid (TUDCA) and trehalose, and the representative HDAC inhibitor VPA (Supplemental Fig. S5A). Treatment with TUDCA and trehalose did not affect *Stat5b* expression in either siControl or siDerl1 NSCs (Supplemental Fig. S5B,C). In contrast, *Stat5b* expression was significantly increased by VPA treatment but not by valpromide (VPM), a VPA analog with no inhibitory effect on HDAC (Supplemental Fig. S5D,E). Consistent with the results for *Stat5b* expression, the percentage of proliferating siDerl1 NSCs were reduced by treatment with VPA but not TUDCA (Supplemental Fig. S5F‒H). These results suggest that the HDAC inhibitor activity of 4-PBA contributes to increased *Stat5b* expression and thus may rescue the impaired transition of Derlin-1- deficient NSCs from active to quiescent states.

**Figure 5.**
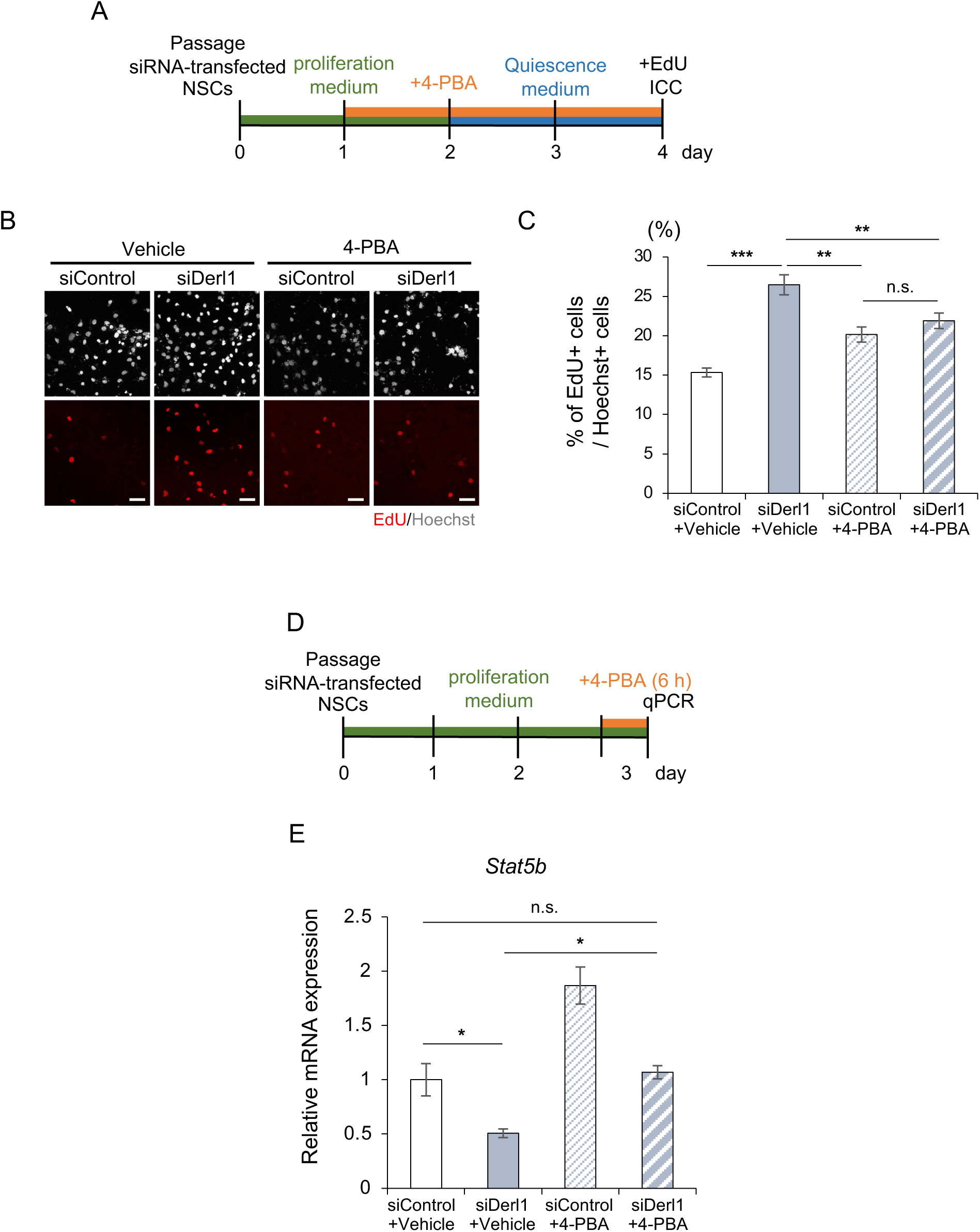
4-PBA induces *Stat5b* expression in NSCs and restores the impaired transition of Derlin-1-deficient NSCs from active to quiescent states. (A) Experimental scheme for evaluating the effect of 4-PBA on the impairment of NSC transition to quiescence by *Derl1* knockdown. (B) Representative images of Hoechst (gray) and EdU (red) staining in siControl and siDerl1 NSCs treated with or without 4-PBA (1 mM). NSCs were fixed 30 min after the addition of EdU. Scale bars: 50 μm. (C) Quantification of the percentage of EdU+ proliferating NSCs among total Hoechst+ cells in 4- PBA-treated siControl and siDerl1 NSCs induced to enter the quiescent state by the administration of BMP4 for 2 days (n = 3; Vehicle, n = 4; 4-PBA). (D) Experimental scheme for assessing the expression of *Stat5b* in siControl and siDerl1 NSCs treated with or without 4-PBA. (E) Expression of *Stat5b* in siControl and siDerl1 NSCs with or without 4-PBA (1 mM) treatment. Gene expression levels were estimated by qPCR and normalized to that of *β-actin* (n = 5; Vehicle, n = 4; 4-PBA). Bar graphs are presented as the mean ± SEM. *P < 0.05, **P < 0.01, and ***P < 0.001 by one-way ANOVA followed by Tukey’s test. n.s., not significant.

### Amelioration of abnormal adult neurogenesis and associated pathological phenotypes by 4-PBA treatment in Derlin-1-deficient mice

4-PBA can easily cross the blood–brain barrier and has been confirmed to exert a therapeutic effect in mouse models of neurological diseases such as Alzheimer’s disease and ALS (Ryu et al. 2005; Wiley et al. 2011). We have reported that 4-PBA administration improved brain atrophy and motor dysfunction in *Derl1^NesCre^*mice (Sugiyama et al. 2022). Therefore, we administered 4-PBA intraperitoneally to *Derl1 ^f/f^* and *Derl1^NesCre^* mice for 14 days and examined its effect on adult neurogenesis *in vivo* (Fig. 6A). Two weeks of 4-PBA administration ameliorated the abnormally increased proliferation of NSCs and the ectopic localization of immature neurons in *Derl1^NesCre^*mice (Fig. 6B,C). When mice intraperitoneally injected with 4-PBA were subjected to the KA-induced seizure susceptibility test (Fig. 6D), 4-PBA treatment alleviated the elevated seizure score in *Derl1^NesCre^* mice (Fig. 6E).

**Figure 6.**
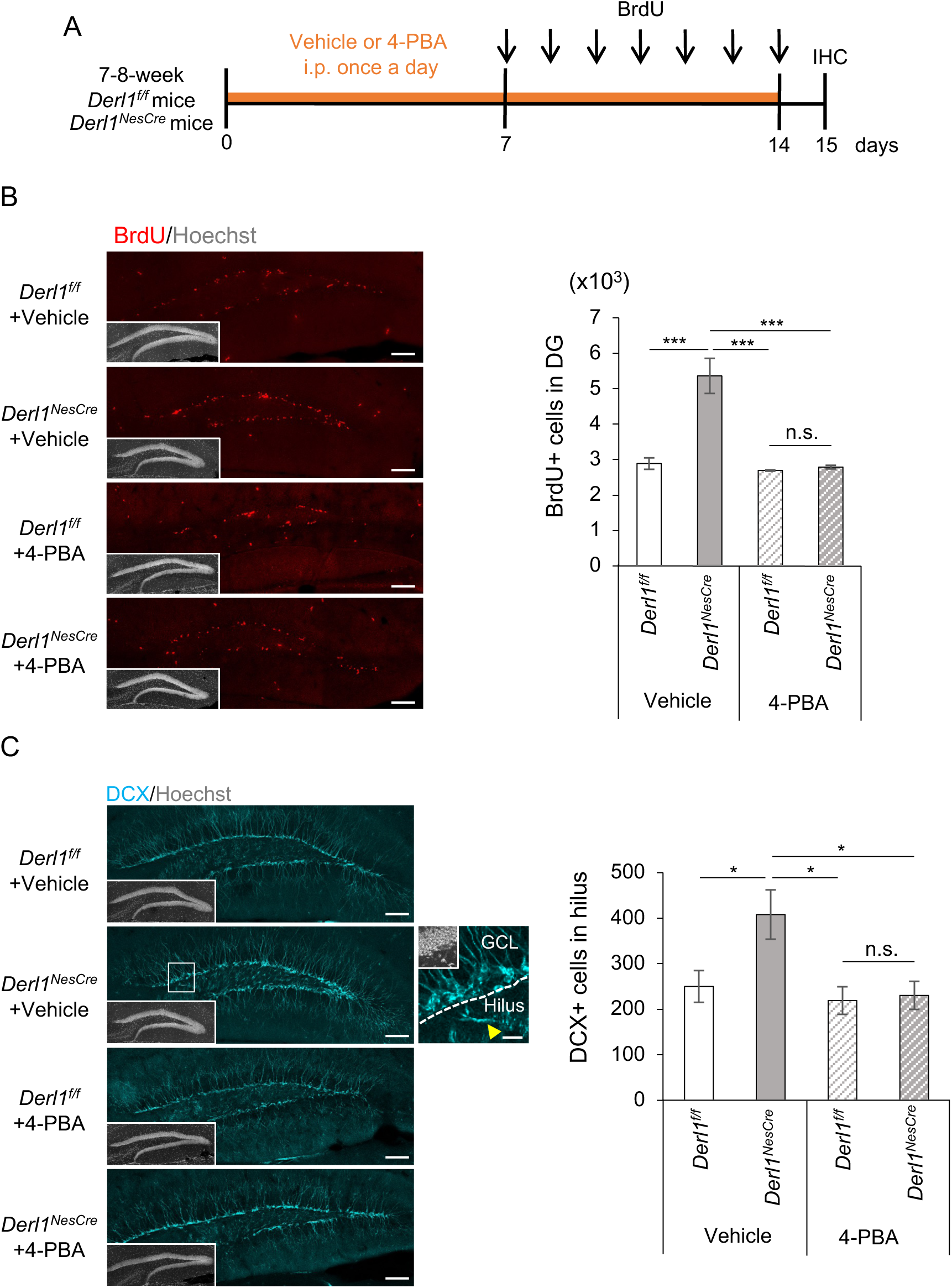
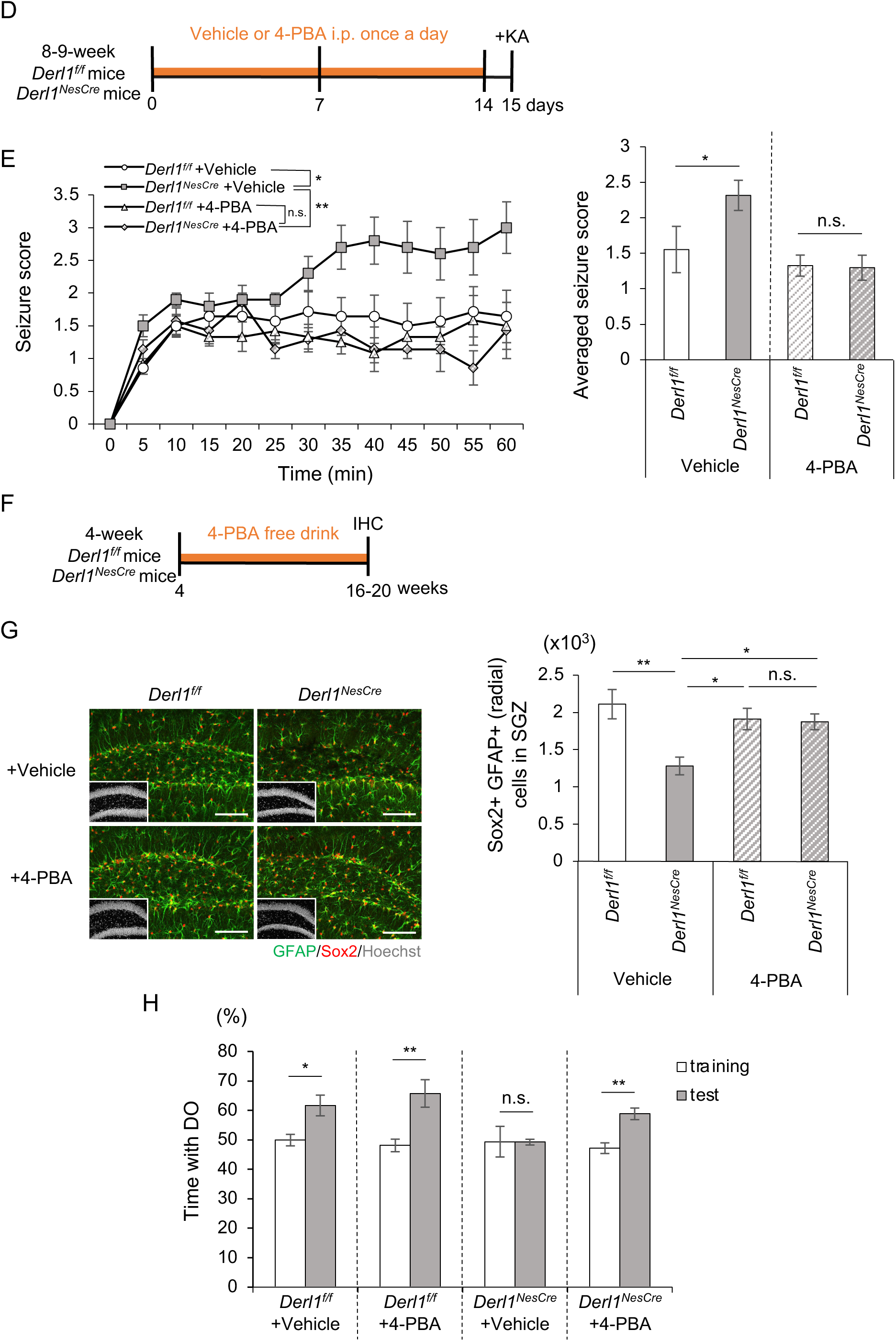
4-PBA improves the increased seizure susceptibility and cognitive dysfunction in *Derl1^NesCre^* mice. (A) Experimental scheme for investigating the proliferation of NS/PCs in *Derl1^f/f^* and *Derl1^NesCre^* mice with or without 4-PBA treatment. *Derl1^f/f^* and *Derl1^NesCre^* mice treated with vehicle or 4-PBA daily for 2 weeks were simultaneously injected with BrdU daily for 7 days during the latter and fixed 1 day after the last BrdU injection. (B) Representative immunofluorescence images of the DG with BrdU (red) and Hoechst staining (gray; insets) (left) and quantification of the number of BrdU+ proliferating cells (right) in the DG of *Derl1^f/f^* and *Derl1^NesCre^* mice treated with or without 4-PBA (n = 3; + Vehicle mice, n = 4; *Derl1^f/f^* + 4-PBA mice, n = 6; *Derl1^NesCre^* + 4-PBA mice). Scale bars: 100 μm. (C) Representative immunofluorescence images of the DG with DCX (cyan) and Hoechst staining (gray; insets). The areas outlined by a white rectangle in the *Derl1^NesCre^* + vehicle panel are enlarged to the right. The yellow arrowhead indicates DCX+ ectopic immature neurons in the hilus, and dashed white lines indicate the boundaries between the GCL and hilus. Scale bars, 100 μm (left images) and 20 μm (right image) (left). Quantification of the number of DCX-positive cells in the hilus in *Derl1^f/f^* and *Derl1^NesCre^* mice treated with or without 4-PBA (right) (n = 6; *Derl1^f/f^* + Vehicle mice, n = 5; *Derl1^NesCre^* + Vehicle mice and *Derl1^NesCre^* + 4-PBA mice, n = 4; *Derl1^NesCre^* + Vehicle mice). (D) Experimental scheme for investigating seizure susceptibility in *Derl1^f/f^* and *Derl1^NesCre^* mice treated with or without 4-PBA. (E) Time plot showing the mean seizure score over 1 h after KA treatment (left) and a bar graph showing the averaged seizure score (right) in *Derl1^f/f^* and *Derl1^NesCre^* mice treated with or without 4-PBA (n = 14; *Derl1^f/f^* + Vehicle mice, n = 10; *Derl1^NesCre^* + Vehicle mice, n = 12; *Derl1 ^f/f^* + 4- PBA mice and n = 7; *Derl1^NesCre^*+ 4-PBA mice). (F) Experimental scheme for assessing the effect of 4-PBA on the depletion of NSCs and cognitive function in *Derl1^f/f^* and *Derl1^NesCre^* mice. 4-PBA solutions were administered from 4 weeks to 16- 20 weeks (4 months) of age through the water supply, which was available ad libitum. (G) Representative immunofluorescence images of the DG with GFAP (green), Sox2 (red), and Hoechst staining (gray; insets) and quantification of the number of Sox2+ radial GFAP+ NSCs in the SGZ of 4-month-old *Derl1^f/f^*and *Derl1^NesCre^* mice treated with or without 4-PBA (n = 5; + Vehicle mice, n = 7; + 4-PBA mice). Scale bars: 100 μm. (H) Percentage of time spent with the displaced object (DO) during the training and testing phases in 4-month-old *Derl1^f/f^* and *Derl1^NesCre^* mice treated with or without 4-PBA (n = 3; + Vehicle mice, n = 5; + 4-PBA mice). Bar graphs are presented as the mean ± SEM. *P < 0.05, **P < 0.01, and ***P < 0.001 by one-way ANOVA followed by Tukey’s test (*B*, *C* and *G*), two-way repeated-measures ANOVA [*E* (left)] or Student’s t test [*E* (right) and *H*]. n.s., not significant.

We next assessed the effect of 4-PBA on NSC depletion and cognitive dysfunction in aged *Derl1^NesCre^*mice. Because long-term intraperitoneal injection stresses the mice, 4-PBA was administered to *Derl1^f/f^* and *Derl1^NesCre^* mice in their ad libitum water supply from 4 weeks to 16-20 weeks of age (Fig. 6F). The number of NSCs in the DG of 4-PBA-treated *Derl1^NesCre^* mice recovered to the same level as that in 4-PBA-treated *Derl1^f/f^* mice, suggesting that the aging-dependent depletion of Derlin-1-deficient NSCs is ameliorated by 4-PBA treatment (Fig. 6G). To examine the effect of 4- PBA treatment on the impaired hippocampus-dependent cognitive function of *Derl1^NesCre^* mice, we performed the novel location recognition test and found that the preference for the DO in the testing phase was restored in 4-PBA-treated *Derl1^NesCre^* mice, as observed in *Derl1^f/f^* mice (Fig. 6H). Our results indicate that the impairment of adult neurogenesis caused by Derlin-1 deficiency and the associated pathological phenotypes, i.e., increased seizure susceptibility and cognitive dysfunction, can be ameliorated by the administration of 4-PBA.

## Discussion

In the present study, we show that Derlin-1 is required for the proper proliferation of NSCs and localization of newborn neurons in the DG through the expression of Stat5b and for brain functions associated with adult hippocampal neurogenesis, including seizure suppression and cognitive function. Furthermore, 4-PBA was found to be effective at rescuing the detrimental phenotypes of Derlin-1- deficient mice via HDAC inhibition.

Maintaining a delicate balance between the quiescent and active states of NSCs is crucial in the adult mammalian brain to prevent depletion and ensure a continuous generation of an adequate number of neurons throughout life (Bond et al. 2015; Shin et al. 2015). Our finding that decreased Stat5b expression increases the number of activated NSCs is consistent with previous reports that Stat5b is required for the maintenance of quiescence in tissue stem cells such as hematopoietic stem cells and hair follicle stem cells (Wang et al. 2009; Wang et al. 2019). A recent study reported that signaling mediated by Stat5 family proteins in the brain is crucial for modulating learning and memory formation but is not associated with adult neurogenesis (Furigo et al. 2018). To clarify the difference between this report and our findings, detailed analysis is required. In addition to decreased expression of Stat5b, other mechanisms driven by Derlin-1 deficiency may also be participated in adult neurogenesis. The tyrosine phosphorylation involved in the transcriptional activation of Stat5b was not required to rescue the abnormal proliferation of Derlin-1-deficient NSCs, and downstream target genes of Stat5b were not enriched among the downstream target genes of Derlin-1. Thus, it is conceivable that the transcriptional activity of Stat5b may not be required to maintain the quiescent state of NSCs. Previous studies have shown that Stat5b, in addition to its role as a transcription factor, is localized to the ER in smooth muscle cells and human pulmonary arterial endothelial cells and is important for maintaining ER structure and mitochondrial function as a nongenomic effect (Lee et al. 2012; Lee et al. 2013; Sehgal 2013). Additionally, Stat5-family proteins without tyrosine phosphorylation are localized in the nucleus and are known to be involved in cytokine-induced megakaryocyte differentiation (Park et al. 2016). Although it is possible that Stat5b may act as a phosphorylation-independent transcription factor or a transcriptional activity–independent factor in the maintenance of NSCs, the Derlin-1-Stat5b axis is a novel and indispensable pathway in adult neurogenesis. Another important issue that remains to be clarified is how Derlin-1 transcriptionally regulates Stat5b expression. Derlin-1 is a well-known ER membrane protein that is indispensable for ER quality control and has no transcriptional activity. The most likely possibility is that the UPR caused by Derlin-1 deficiency inhibits the expression of *Stat5b* mRNA in the activated NSCs. Although ER stress was also induced in the brains of Derlin-2-deficient mice (Supplemental Fig. S1I) (Sugiyama et al. 2021), abnormal proliferation of NSCs was not observed in Derlin-2-deficient mice (Fig. S1*J* and *K*). The possible involvement of ER stress cannot be ruled out, but it is strongly suggested that Derlin-1-specific downstream targets may contribute to the transcriptional regulation of *Stat5b*. Further studies are important to clarify how Stat5b expression is regulated by Derlin-1; such studies will provide novel insight into the mechanism of adult neurogenesis.

Derlin-1 deficiency induces the ectopic localization of newborn neurons in the hilus, which is due to abnormal migration. Among the factors involved in cell migration, the expression of CXC motif chemokine receptor 4 (Cxcr4), which is indispensable in NS/PCs, has been implicated in the appropriate localization of newborn neurons in the adult DG (Schultheiss et al. 2013; Sakai et al. 2018). Although *Cxcr4* was not found in differentially expressed genes in siDerl1 NSCs in our RNA-seq analysis (Supplemental Table S1), it may be possible that the expression of Cxcr4 protein on the plasma membrane surface is suppressed by Derlin-1 deficiency. It is also possible that abnormally proliferated NSCs or newborn neurons might be physically extruded from the DG to the hilus or that Stat5b directly or indirectly regulates the location of adult neurogenesis. The mechanism by which the survival of NS/PCs, which proliferate in the adult DG, is ultimately reduced in Derlin-1-deficient mice is also unknown. Although further research is needed to elucidate these unresolved mechanisms, our results indicate that Derlin-1 regulates adult neurogenesis in a spatiotemporal manner.

ER stress is thought to be involved in the pathogenesis of neurological diseases, including ALS and spinocerebellar ataxia (Nishitoh et al. 2002; Nishitoh et al. 2008; Ghemrawi and Khair 2020). Chemical chaperone therapy is currently being developed as a treatment for several diseases, including some neurological diseases, with the aim of reducing ER stress. For example, clinical trials are currently being conducted on ALS patients using sodium phenylbutyrate (a salt of 4-PBA) and TUDCA, which have shown effects such as delayed disease progression and prolonged survival (Paganoni et al. 2020; Paganoni et al. 2022). However, it is questionable whether 4-PBA improves the pathology of neurological disease through its chaperone activity alone. Other chemical chaperones, TUDCA and trehalose, had no effect on *Stat5b* expression, while VPA increased its expression, suggesting that HDAC inhibition by 4-PBA may function to restore *Stat5b* expression in Derlin-1- deficient NSCs. Since both 4-PBA and VPA are short-chain fatty acid group HDAC inhibitors that mainly inhibit class I HDACs (HDAC1, 2, 3 and 8) (de Ruijter et al. 2003), it is possible that activation of class I HDACs in NSCs may suppress *Stat5b* expression. VPA is known to inhibit NS/PCs proliferation by inducing cyclin-dependent kinase inhibitors p21 through its HDAC inhibitor activity (Jessberger et al. 2007). In this study, we discovered a novel function of the HDAC inhibitor 4-PBA in regulating adult neurogenesis by inducing specific genes, including *Stat5b*. Although further studies are needed to elucidate the precise molecular mechanisms by which HDAC inhibitors ameliorate abnormal adult neurogenesis in Derlin-1-deficient mice, this study demonstrates that the administration of HDAC inhibitors such as 4-PBA and VPA may be applicable in research aiming to clarify the pathological mechanisms of diseases caused by the disruption of adult neurogenesis.

In summary, the Derlin-1-Stat5b axis is essential for maintaining adult neurogenesis throughout life. Maintenance of adult neurogenesis via Derlin-1 function is essential for controlling seizure susceptibility and maintaining cognitive function, and pathologies caused by its disruption are ameliorated by HDAC inhibition. Our discovery paves the way for the elucidation of mechanisms and the possible treatment of neurological diseases caused by abnormal adult neurogenesis, depending on further research.

## Materials and methods

### Animals

All mice used in this experiment were raised under specific-pathogen-free conditions, housed under a 12-h/12-h light/dark cycle, and fed ad libitum. Details regarding *Derl1^f/f^* mice, *Derl2^f/f^* mice, and mice expressing Cre recombinase driven by the *nestin* or *CaMKIIa* promoter have been described in previous reports (Isaka et al. 1999; Dougan et al. 2011; Karpati et al. 2019; Sugiyama et al. 2021). These mice were intercrossed to generate *Derl1^NesCre^* mice, *Derl1^CaMKIIαCre^* mice and *Derl2^NesCre^* mice. Both male and female mice were used. All mouse experiments were approved by the Animal Research Committee of the University of Miyazaki following institutional guidelines. The experiments were conducted according to institutional guidelines. All efforts were made to minimize animal suffering and reduce the number of animals used.

### Cell lines

Human embryonic kidney (HEK) 293T cells were obtained from the American Type Culture Collection (ATCC). HEK293T cells were grown in DMEM (08459-64, Nacalai Tesque) supplemented with 10% FBS and penicillin‒streptomycin solution (09367-34, Nacalai Tesque). Adult hippocampal NSCs were isolated and cloned from Fisher 344 rats and characterized in previous reports (Palmer et al. 1997; Mira et al. 2010). Adult hippocampal NSCs were cultured in DMEM/F12 supplemented with N2, penicillin‒streptomycin solution, and bFGF (20 ng/mL) (100-18B, PeproTech) (proliferation medium) or bFGF (10 ng/mL) and diazepam (100 μM) (045-18901, Wako) or BMP4 (50 ng/mL) (5020-BP, R&D Systems) (quiescence medium) on coated culture dishes with poly-L-ornithine (P- 3655, Sigma‒Aldrich) and laminin (354232, Corning). All cells were maintained under a 5% CO2 atmosphere at 37°C.

### siRNA transfection

siRNA transfection was performed using Lipofectamine RNAiMAX reagent (56532, Invitrogen). The following siRNAs were used for knockdown of adult rat-derived hippocampal NSCs: Stealth RNAi™ siRNA Derl1-MSS289837 (Invitrogen), Stealth RNAi™ siRNA Stat5b-RSS332572 (Invitrogen). Stealth RNAi™ siRNA Negative Control Med GC Duplex (452001, Invitrigen) was used as the control.

### BrdU administration

To label proliferating cells, BrdU (B5002, Sigma‒Aldrich) dissolved in saline (0.9% NaCl) was injected (50 mg/kg) intraperitoneally into 8- or 9-week-old mice once a day for 7 days. The mice were sacrificed 1 day or 3 weeks after the last BrdU injection.

### Tissue preparation for immunofluorescence

Mice were deeply anesthetized by intraperitoneal injection of a 4 mg/kg midazolam/0.3 mg/kg medetomidine/5 mg/kg butorphanol mixture and transcardially perfused with phosphate-buffered saline (PBS) followed by 4% paraformaldehyde (PFA) in PBS. Brains were dissected and postfixed overnight in 4% PFA at 4°C. Fixed brains were incubated in 15% sucrose solution at 4°C overnight, followed by 30% solution at 4°C overnight. Brains were then cut into two pieces along the midline, and each half was embedded in an optimal cutting temperature compound (4583, Tissue Tek; Sakura Finetek) and stored at −80°C. Embedded frozen brains were serially sectioned in the coronal plane at 40-μm thickness using a freezing microtome (CM3050S, Leica Microsystems). Every sixth section was sequentially transferred to 6-well plates of PBS for subsequent immunohistochemical staining.

### Immunohistochemistry

The bran sections were washed with PBS and incubated in blocking buffer (PBS containing 3% FBS and 0.1% Triton X-100) for 1 h at room temperature (RT), followed by overnight incubation at 4°C with the primary antibody diluted in blocking buffer. Sections were washed three times with PBS and incubated for 2 h at RT with a secondary antibody diluted in blocking buffer. After a third wash with PBS, the sections were mounted on glass slides with Immu-Mount (9990402, Thermo Scientific). For the staining of BrdU, sections were incubated with 2 N HCl at 37°C for 15 min and washed with PBS 3 times before being blocked. Immunofluorescence images were acquired using a confocal laser microscope (TSC-SP8, Leica Microsystems) and processed using Adobe Photoshop Elements 2021 (Adobe). Nuclei were counterstained using bisbenzimide H33258 fluorochrome trihydrochloride solution (Hoechst; 19173-41, Nacalai Tesque). Antibodies are listed in the Supplemental Table S2.

### Cell counting in brain sections

Quantifying the number of respective marker-positive cells in the DG was performed using every sixth hemisphere section. The number of marker-positive cells was counted and multiplied by 6 to estimate the total number of DGs. A cell was determined to be located in the hilus if the soma of the cell was clearly located on the hilus side relative to the continuous line drawn between the SGZ and the hilus.

### In vitro cell proliferation assay

To label proliferating NSCs, 10 mg/mL EdU from a Click-iT EdU Alexa Fluor 555 Imaging Kit (C10338, Invitrogen) was added to the culture medium 30 min before fixation. EdU staining was performed following the kit manufacturer’s instructions, followed by immunocytochemistry (below).

### Immunocytochemistry

Adult hippocampal NSCs were fixed with 4% PFA in PBS for 20 min, washed three times in PBS after EdU staining, permeabilized, blocked with blocking buffer (PBS containing 3% FBS and 0.1% Triton X-100) for 30 min at RT, and incubated for 1.5 h at RT with the indicated primary antibody diluted in blocking buffer. Cells were washed three times with PBS and incubated for 1.5 h at RT with the secondary antibody diluted in blocking buffer. After a third wash with PBS, cells were mounted with Immu-Mount (Thermo Scientific) on glass slides. Nuclei were counterstained using Hoechst (1:500; Nacalai Tesque). Immunofluorescence images were obtained using a confocal laser microscope (Leica Microsystems) and processed using Adobe Photoshop Elements 2021. The antibodies are listed in the Supplemental Table S2.

### Immunoblotting

The adult hippocampal NSCs were lysed in lysis buffer (20 mM Tris-HCl pH 7.5, 150 mM NaCl, 5 mM EGTA, and 1% Triton X-100) supplemented with 5 μg/mL leupeptin (43449-62, Nacalai Tesque). These whole-cell lysates were resolved by sodium dodecyl sulfate‒polyacrylamide gel electrophoresis (SDS‒PAGE) and blotted onto polyvinylidene fluoride (PVDF) membranes. After blocking with 5% skim milk in TBS-T (50 mM Tris-HCl pH 8.0, 150 mM NaCl, and 0.05% Tween-20), the membranes were probed with the indicated antibodies, and immunolabeling was detected using an enhanced chemiluminescence (ECL) system. The antibodies are listed in the Supplemental Table S2.

### Lentivirus production

Lentiviruses were produced by co-transfecting HEK293T cells with the lentivirus constructs pRRL- Venus-HA, pRRL-Stat5b-Venus, pRRL-Stat5b-HA or pRRL-Stat5b (Y699F)-Venus, and lentivirus packaging vector constructs pMD2.G (12259, Addgene) and psPAX2 (12260, Addgene) using Polyethylenimine (PEI)-Max (24765-1, Polysciences). The culture medium was changed at 16–24 h after transfection. The supernatants were collected at 24 and 48 h after a medium change and centrifuged at 6,000 x g overnight at 4°C. After discarding the supernatant, the virus solution was resuspended in 1 mL of new medium per 10 cm dish (*in vitro* conditions), and then the virus solution was concentrated using the Lenti-X Concentrator (631231, Clontech) and suspended in PBS (*in vivo* conditions).

### In vitro lentiviral infection

The virus solutions were introduced into adult hippocampal NSCs by adding these supernatants to the culture 24 h after passaging. At 48 h after infection, the medium was replaced with quiescence medium. The cells were cultured for another 48 h and then fixed for an EdU-labeled cell proliferation assay.

### In vivo lentiviral infection

Nine-week-old *Derl1^NesCre^* mice were anesthetized by intraperitoneal injection of a 4 mg/kg midazolam/0.3 mg/kg medetomidine/5 mg/kg butorphanol mixture. The virus suspension was injected stereotaxically into the bilateral DG using the following coordinates relative to bregma: caudal, −2.0 mm; lateral, ±1.5 mm; ventral, −2.3 mm. In each DG, 1.5 µL of lentivirus was injected over 1 min using a 5 µL Hamilton syringe. Two weeks after the lentiviral injection, the brains were fixed for immunohistochemistry. Mice lacking HA-tag-positive cells in the DG were excluded from the study.

### Quantitative real-time PCR analysis

Total RNA was isolated from adult hippocampal NSCs using RNAiso Plus (9109, Takara Bio) and reverse transcribed using RevaTra Ace qPCR RT Master Mix with gDNA Remover (FSQ-301, TOYOBO) according to the manufacturer’s instructions. Quantitative PCR was performed using SYBR Green PCR Master Mix (KK4602, Kapa Biosystems) and a StepOnePlus Real-Time PCR System (Applied Biosystems). Expression levels were normalized to the expression of *β-actin* mRNA and calculated relative to the control. The following primers were used for quantitative PCR: β-actin primer forward 5′-TCCTCCCTGGAGAAGAGCTAT-3′, β-actin primer reverse 5′- TCCTGCTTGCTGATCCACAT-3′, Stat5b primer forward 5′-GGGCATCACCATTGCTTGGAAG-3′, Stat5b primer reverse 5′- CCGGATAGAGAAGTCTCTTGTGG-3′

### DNA microarray analysis, RNA-seq

Standard procedures were used for DNA microarray and RNA-seq analyses; specific details are documented in the Supplemental Material. Data generated in this study have been deposited in GEO under accession number GSE226345 and GSE229251.

### Seizure behavioral assays

Seizures were induced in 8- to 12-week-old *Derl1^f/f^* and *Derl1^NesCre^* mice by intraperitoneal injections of 20 mg/kg KA (BML-EA123, Enzo Life Sciences) dissolved in distilled H2O. The behavior of the mice was observed for 1 h after the injection, and a seizure score was recorded manually every 5 min. The seizure score was modified into five stages from the previously described criteria (Racine 1972). Briefly, the following seizure scale was used: normal exploratory activity (0), staring and reduced locomotion (1), immobility with fast breathing/scratching behavior (2), repetitive head and limb movements (3), sustained rearing with forelimb clonus (4), and full body extension (full tonic extension) and death (5).

### Novel location recognition test

*Derl1^f/f^* and *Derl1^NesCre^* mice were placed in a white plastic chamber (45 × 45 × 43 [H] cm) that contained two identical objects in adjacent corners; the mice were allowed to explore the objects freely for 3 min and then taken back to their home cage for 3 min, completing one training session. After three repetitions of the training session, one of the objects was moved to the opposite side of the corner of the chamber and allowed to freely explore the familiar and displaced objects for 15 min (Testing session). All sessions were recorded with an overhead video, and exploration behavior was defined as activities such as sniffing and rearing against the object. The time spent exploring each object during the training and test sessions was scored manually. The exploration ratio for objects in novel locations (displaced objects) was calculated using the formula t displaced /(t displaced + t familiar), as described previously (Mumby et al. 2002). The chamber and objects were cleaned with 70% ethanol before the next mouse was tested.

### 4-PBA administration

Intraperitoneal injections of 200 mg/kg 4-PBA (820986, MERCK or P21005, Sigma‒Aldrich) were performed daily from 8-9 to 10-11 weeks of age for immunohistochemistry and seizure behavioral assays. To examine the depletion of NSCs and cognitive function, 10 mg/mL of 4-PBA (Sigma‒ Aldrich) solution was administered in the ad libitum water supply from 4 weeks to 16-20 weeks of age.

### Statistical analysis

All data are presented as the means ± standard errors. Student’s t test was performed to compare two group means. One-way ANOVA followed by post hoc tests compared three or more group means. Data from the 1 h trial of seizure behavior were analyzed by two-way repeated-measures ANOVA, and post hoc analysis was performed using Bonferroni’s multiple comparison test. Statistical analyses were performed using EZR software version 1.30 (Kanda 2013) or GraphPad Prism 9 (GraphPad Software). A *P <* 0.05 (two-tailed) was considered significant for all tests.

## Acknowledgements

We thank Dr. R. Kageyama (RIKEN) for *Tg(Nes-Cre)1Kag* mice, Dr. Ploegh HL (Boston Children’s Hospital and Harvard Medical School) for *Derl2^f/f^* mice, Drs. A. Futatsugi (Kobe City College of Nursing) and K Mikoshiba (RIKEN, Shanghai Tech University and Toho University) for *TgN(a-CaMKII-nlCre)/10* mice, Dr. D. Trono (Ecole Polytechnique Fédérale de Lausanne) for lentivirus packaging vector constructs, and Dr. K. Takao (University of Toyama) for technical assistance and discussion. This study was supported by MEXT/JSPS KAKENHI (grant number, 22K06254 [N.M.], 21K06175 [H.K.], 23H00391 [K.N.], 18H02973 and 22H02954 [H.N.]), AMED (grant number JP22gm6410024 [H.K.] and JP20gm1310008 [K.N.]), Terumo Life Science Foundation (H.N.), Naito Foundation (H.N.), Ono Medical Research Foundation (N.M.), Takeda Science Foundation (H.N.), TMDU Nanken-Kyoten Grant Number 2022-39 (H.N.), and Joint Usage and Joint Research Programs, Institute of Advanced Medical Sciences, Tokushima University (N.M., H.N.).

## Author contributions

N.M. conceptualization, investigation, funding acquisition, and writing-original draft. T.M. investigation. H.K. conceptualization, funding acquisition, supervision, and writing-review and editing. Y.M., and K.T. investigation and methodology. T.K. resources and methodology. K.N. resources, supervision and writing-original draft, review, and editing. H.N. conceptualization, data curation, supervision, funding acquisition, project administration and writing-original draft, review, and editing.

