## Supplemental Information for "Homeostatic Regulation of Seizure Susceptibility and Cognitive Function by Derlin-1 through Maintenance of Adult Neurogenesis"

[1]Laboratory of Biochemistry and Molecular Biology, Department of Medical Sciences, University of Miyazaki, 889-1692 Miyazaki, Japan
[2]Department of Stem Cell Biology and Medicine, Graduate School of Medical Sciences, Kyushu University, 812-8582 Fukuoka, Japan
[3]Division of Genome Medicine, Tokushima University, 770-8503 Tokushima, Japan
[4]Genome Laboratory, Medical Research Institute, Tokyo Medical and Dental University, 113-8510 Tokyo, Japan
[5]Frontier Science Research Center, University of Miyazaki, 889-1692 Miyazaki, Japan
[6]Lead contact

**This file includes:**

Supplemental Figure 1; related to Figure 1
Supplemental Figure 2; related to Figure 2
Supplemental Figure 3; related to Figure 3
Supplemental Figure 4; related to Figure 4
Supplemental Figure 5; related to Figure 5
Supplemental Table S1

Supplemental Material and Methods
SI References
Supplemental Table S2

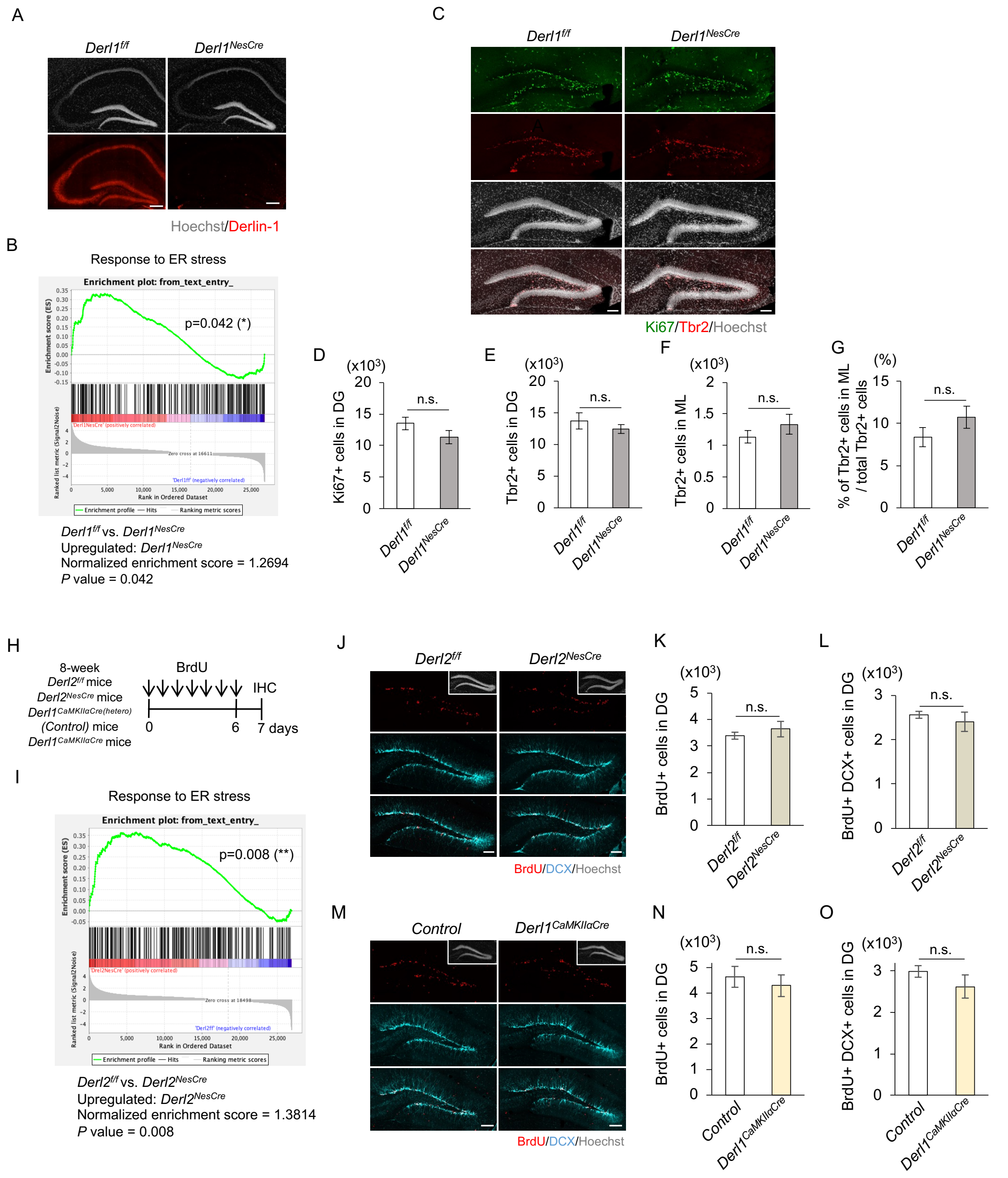


**Supplemental Figure 1. Loss of *Derl1*, but not *Derl2*, specifically promotes NSC activation in the adult DG.**

(*A*) Representative immunofluorescence images with Hoechst (gray) and Derlin-1 staining (red) in the adult hippocampus of *Derl1*$^{f/f}$ and *Derl1*$^{NesCre}$ mice. Scale bar: 100 µm.

(*B*) GSEA showing differential expression of 238 genes in the DG categorized by the GO term "Response to ER stress." GSEA shows gene expression changes in the DG of *Derl1*$^{NesCre}$ mice relative to *Derl1*$^{f/f}$ mice. The enrichment plot shows the distribution of genes in each set that are positively (red) or negatively (blue) correlated with Derlin-1 deficiency.

(*C*) Representative immunofluorescence images with Ki67 (green), Tbr2 (red), and Hoechst (gray) staining of the DG in postnatal day 14 (P14) *Derl1*$^{f/f}$ and *Derl1*$^{NesCre}$ mice. Scale bars: 100 µm.

(*D–F*) Quantification of the numbers of Ki67+ cells (*D*) and Tbr2+ cells (*E*) in the DG as well as the number of Tbr2+ cells (*F*) in the molecular layer (ML) of the DG (n = 3 mice).

(*G*) The percentage of Tbr2+ cells in ML among total Tbr2+ cells in the DG of P14 *Derl1*$^{f/f}$ and *Derl1*$^{NesCre}$ mice (n = 3 mice).

(*H*) Experimental scheme for investigating the proliferation of NS/PCs and neurogenesis in *Derl2*$^{f/f}$, *Derl2*$^{NesCre}$, *Derl1*$^{CaMKII\alpha Cre(hetero)}$ (Control), and *Derl1*$^{CaMKII\alpha Cre}$ mice.

(*I*) GSEA showing differential expression of 238 genes in the DG categorized by the GO term "Response to ER stress." GSEA shows gene expression changes in the DG of *Derl2*$^{NesCre}$ mice relative to *Derl2*$^{f/f}$ mice. The enrichment plot shows the distribution of genes in each set that are positively (red) or negatively (blue) correlated with Derlin-2 deficiency.

(*J–O*) Representative immunofluorescence images of the DG with BrdU (red), DCX (cyan), and Hoechst staining (gray; insets) (*J* and *M*) and quantification of BrdU+ proliferating cells (*K* and *N*) or BrdU+ DCX+ newborn immature neurons (*L* and *O*) in mice of each genotype (n = 3 mice). Scale bars: 100 μm.

Bar graphs are presented as the mean ± SEM. n.s., not significant.

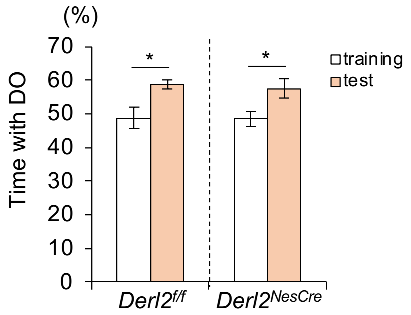


**Supplemental Figure 2. Loss of *Derl2* does not impair cognitive function.**
Percentage of time spent with the displaced object (DO) during the training and testing phase in 4-month-old *Derl2*$^{f/f}$ and *Derl2*$^{NesCre}$ mice (n = 5; *Derl2*$^{f/f}$ mice, n = 3; *Derl2*$^{NesCre}$ mice).
Bar graphs are presented as the mean ± SEM. *P < 0.05 by Student's t test.

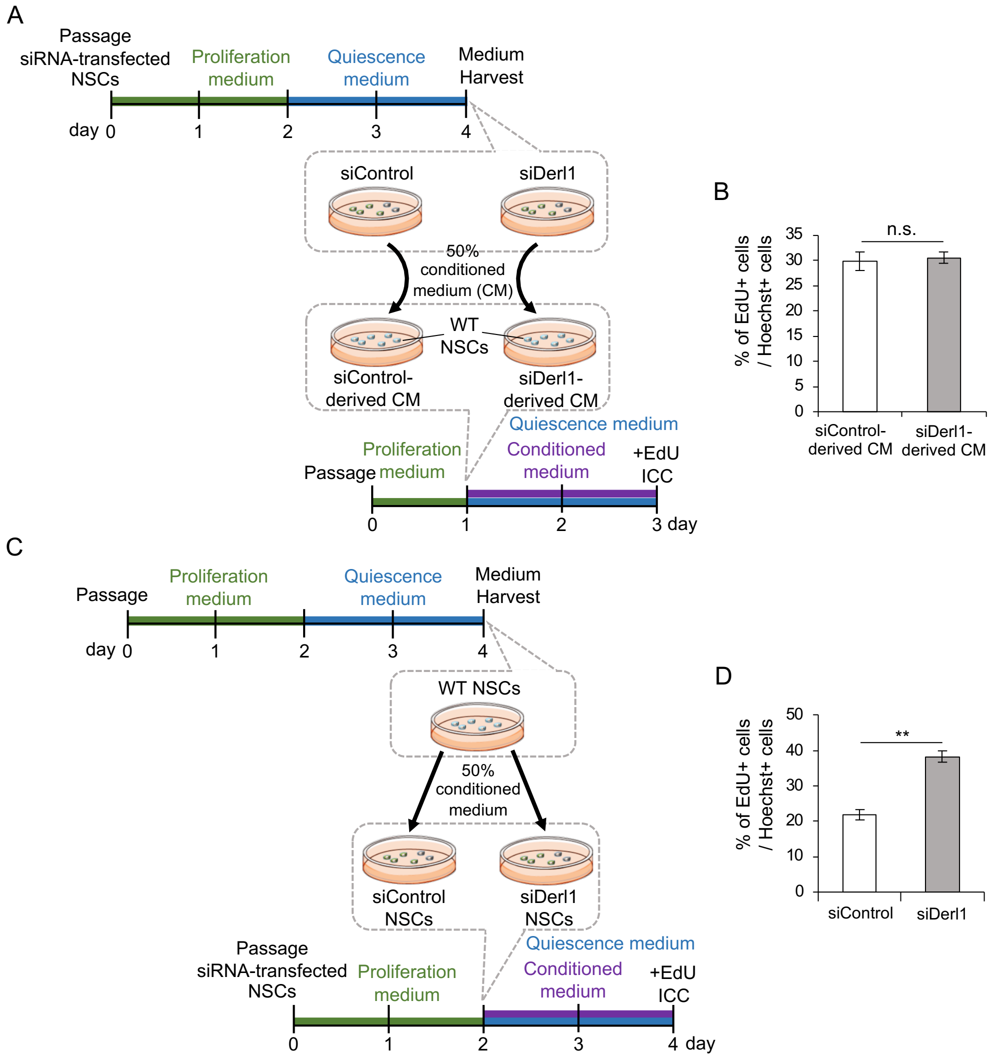


**Supplemental Figure 3. Inhibition of the transition from active to quiescent states in Derlin-1-deficient NSCs is cell-autonomously regulated.**
(*A*) Experimental scheme to investigate NSC proliferation with conditioned medium derived from control and *Derl1* knockdown NSCs over 2 days.
(*B*) Quantification of the percentage of EdU+ proliferating NSCs among total Hoechst+ cells cultured for 2 days in siControl and siDerl1 NSC–derived conditioned quiescence medium (n = 4).
(*C*) Experimental scheme to investigate NSC proliferation of control and *Derl1* knockdown NSCs with conditioned medium derived from WT NSCs over 2 days.
(*D*) Quantification of the percentage of EdU+ proliferating NSCs among total Hoechst+ cells cultured for 2 days in siControl and siDerl1 NSCs with WT NSC–derived conditioned quiescence medium (n = 3).
Bar graphs are presented as the mean ± SEM. **P < 0.01 by Student's t test. n.s., not significant.

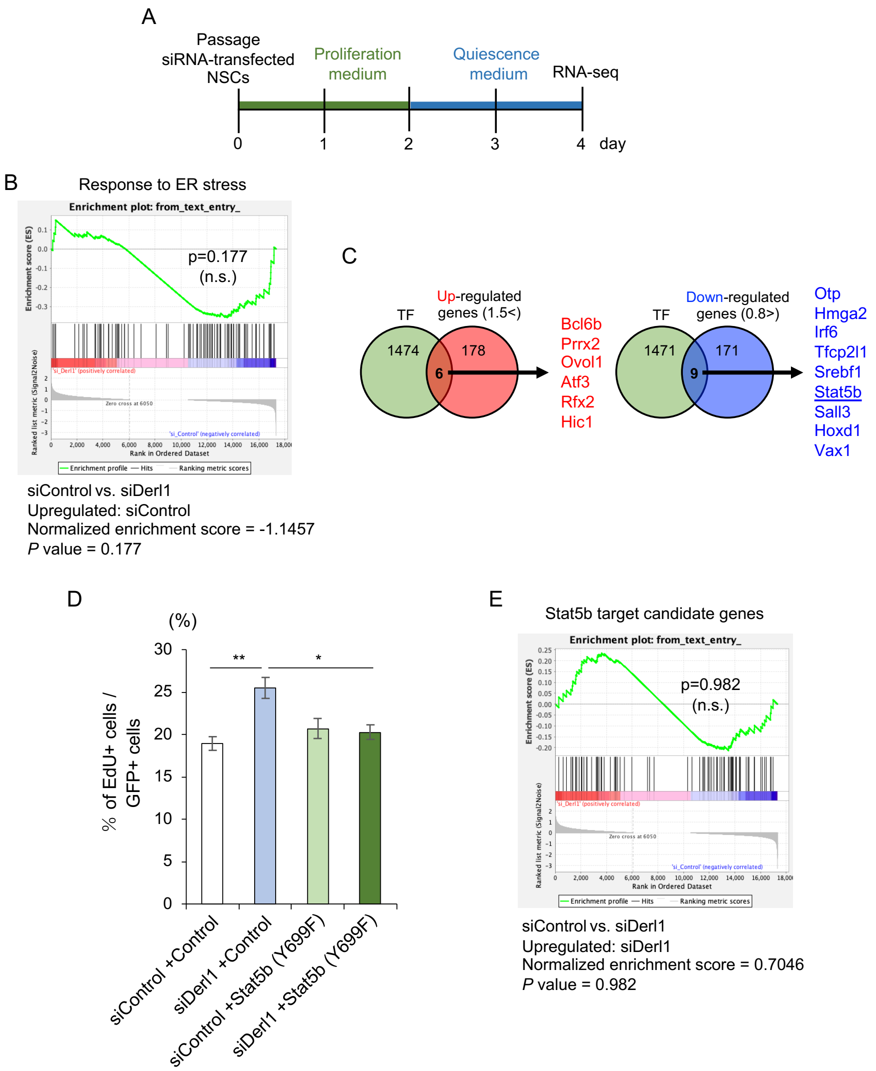


**Supplemental Figure 4. *Stat5b* expression is decreased in Derlin-1-deficient NSCs, and the phosphorylation of Stat5b (Y699) is not required for the rescue of abnormal proliferation of Derlin-1-deficient NSCs.**

(*A*) Experimental scheme for investigating the molecular mechanism underlying the impairment of NSC transition to quiescence by *Derl1* knockdown.

(*B*) GSEA showing differential expression of 92 genes in the NSCs categorized by the GO term "Response to ER stress." GSEA shows gene expression changes in siDerl1 NSCs relative to siControl NSCs. The enrichment plot shows the distribution of genes in each set that are positively (red) or negatively (blue) correlated with *Derl1* knockdown.

(*C*) Venn diagrams showing the overlap between transcription factor (TF) genes and upregulated (left) or downregulated (right) genes in siDerl1 NSCs.

(*D*) Quantification of the percentage of EdU+ proliferating NSCs among total GFP+ cells in siControl and siDerl1 NSCs with or without exogenous expression of mutant Stat5b (Y699F) (n = 3).

(*E*) GSEA showing differential expression of 80 candidate Stat5b target genes. GSEA shows gene expression changes in siDerl1 NSCs relative to siControl NSCs. The enrichment plot shows the distribution of genes in each set that are positively (red) or negatively (blue) correlated with *Derl1* knockdown.

Bar graphs are presented as the mean ± SEM. *P < 0.05 and **P < 0.01 by one-way ANOVA followed by Tukey's test.

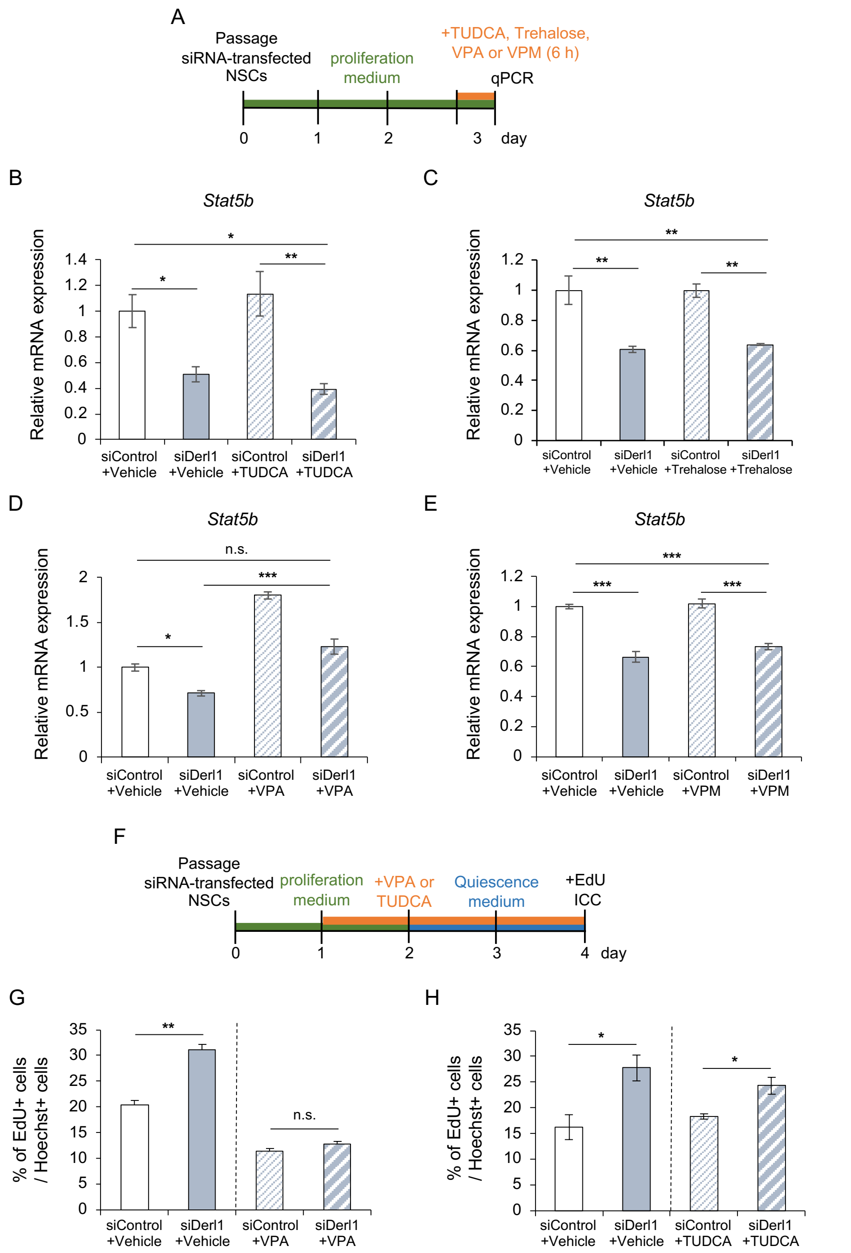


**Supplemental Figure 5. HDAC inhibitory activity, but not chaperone activity, increases *Stat5b* expression and inhibits the proliferation of NSCs.**

(*A*) Experimental scheme for assessing the expression of *Stat5b* in siControl and siDerl1 NSCs treated with or without TUDCA (50 μM), trehalose (10 mM), VPA (1 mM), or VPM (1 mM).

(*B–E*) Expression of *Stat5b* in siControl and siDerl1 NSCs with or without TUDCA (*B*) (n = 4; Vehicle, n = 3; TUDCA), trehalose (*C*) (n = 4), VPA (*D*) (n = 3), or VPM (*E*) treatment (n = 3). Gene expression levels were estimated by qPCR and normalized to that of *β-actin*.

(*F*) Experimental scheme for evaluating the effect of VPA (1 mM) or TUDCA (50 μM) on the impairment of the transition of NSCs to quiescence by *Derl1* knockdown.

(*G*,*H*) Quantification of the percentage of EdU+ proliferating NSCs among total Hoechst+ cells in VPA-treated (*G*) (n = 3) or TUDCA-treated (*H*) (n = 4; Vehicle, n = 5; TUDCA) siControl and siDerl1 NSCs induced to enter the quiescent state by the administration of BMP4 for 2 days.

Bar graphs are presented as the mean ± SEM. *P < 0.05, **P < 0.01, and ***P < 0.001 by one-way ANOVA followed by Tukey's test (*B–E*) or Student's t test (*G* and *H*). n.s., not significant.

### Supplemental Table S1. List of genes with altered expression in siDerl1 NSCs

**Gene list whose expression level is >1.5-fold increased in siDerl1 NSCs**

| Refseq ID | GeneSymbol | Description | Fold change | P value |
|---|---|---|---|---|
| NM_053789 | Il17b | interleukin 17B | 28.72369561 | 0.003845464 |
| NM_017154 | Xdh | xanthine dehydrogenase | 28.01189405 | 2.90E-05 |
| NM_138854 | Slc38a5 | solute carrier family 38, member 5 | 27.30578913 | 0.016246065 |
| NM_001106277 | Saxo2 | stabilizer of axonemal microtubules 2 | 23.00916878 | 0.012429043 |
| NM_001105749 | Il16 | interleukin 16 | 19.66806675 | 0.033194297 |
| NM_001011971 | Xkr4 | XK related 4 | 18.38027887 | 0.03431546 |
| NM_020101 | Adap2 | ArfGAP with dual PH domains 2 | 18.18781236 | 0.026796782 |
| NM_001271361 | St6galnac5 | ST6 N-acetylgalactosaminide alpha-2,6-sialyltransferase 5 | 18.14166765 | 0.046077641 |
| NM_001106906 | Gpr45 | G protein-coupled receptor 45 | 17.83467498 | 0.046662042 |
| NM_001107021 | Hic1 | HIC ZBTB transcriptional repressor 1 | 17.64655338 | 0.0399173 |
| NM_001305243 | Eda | ectodysplasin-A | 17.4413855 | 0.044453758 |
| NM_001108279 | Bcl6b | BCL6B, transcription repressor | 13.01440305 | 0.000679595 |
| NM_130741 | Lcn2 | lipocalin 2 | 12.265538 | 2.40E-05 |
| NM_024354 | Chrna4 | cholinergic receptor nicotinic alpha 4 subunit | 11.6910145 | 0.003002521 |
| NM_001007011 | Zpbp2 | zona pellucida binding protein 2 | 11.25355364 | 0.002292234 |
| NM_001108979 | Atp6v1e2 | ATPase H+ transporting V1 subunit E2 | 9.452958238 | 0.009062017 |
| NM_016994 | C3 | complement C3 | 9.355076006 | 0.000269583 |
| NM_001271398 | Kirrel2 | kirre like nephrin family adhesion molecule 2 | 9.002633066 | 0.006733583 |
| NM_001013084 | Akr1b10 | aldo-keto reductase family 1 member B10 | 8.853624237 | 0.010955202 |
| NM_001044304 | Eid3 | EP300 interacting inhibitor of differentiation 3 | 8.502484989 | 0.001164639 |
| NM_207607 | Ns5atp4 | | 8.199592209 | 0.029474077 |
| NM_194353 | Setl1 | | 8.0750904 | 0.028780778 |
| NM_001033957,N | Kcng3 | potassium voltage-gated channel modifier subfamily G member 3 | 8.062650312 | 0.018451264 |
| NM_031055 | Mmp9 | matrix metallopeptidase 9 | 7.4276006 | 1.55E-06 |
| NM_001109536 | Ptx3 | pentraxin 3 | 7.249857943 | 0.032714643 |
| NM_013107 | Bmp6 | bone morphogenetic protein 6 | 7.167416648 | 0.000135371 |
| NM_001105739 | Prrx2 | paired related homeobox 2 | 6.075331872 | 0.013136782 |
| NM_001108550 | Postn | periostin | 6.012517256 | 0.005051686 |
| NM_001007612 | Ccl7 | C-C motif chemokine ligand 7 | 5.660284185 | 0.005588881 |
| NM_001104527 | Prr15 | | 5.651010713 | 0.004505096 |
| NM_001109996,N | Apoc1 | apolipoprotein C1 | 5.634992027 | 0.020834602 |
| NM_020305 | Adam4 | a disintegrin and metalloprotease domain 4 | 5.245130505 | 0.001996655 |
| NM_022931 | Rims3 | regulating synaptic membrane exocytosis 3 | 4.824446032 | 0.02447858 |
| NM_001106664 | Tyrp1 | tyrosinase-related protein 1 | 4.823241381 | 0.005019777 |
| NM_139216,NM_1 | Kcnc2 | potassium voltage-gated channel subfamily C member 2 | 4.649878817 | 0.006359266 |
| NM_001108161 | Cilp | cartilage intermediate layer protein | 4.609237491 | 0.000349139 |
| NM_030845 | Cxcl1 | C-X-C motif chemokine ligand 1 | 4.450594639 | 0.001632821 |
| NM_001191861 | Cdk6 | cyclin-dependent kinase 6 | 4.304971631 | 0.018957314 |
| NM_153722 | Mrgprf | MAS related GPR family member F | 4.274727183 | 0.011359421 |
| NM_031530 | Ccl2 | C-C motif chemokine ligand 2 | 4.189269844 | 0.000561563 |
| NM_207613 | Cdh15 | cadherin 15 | 4.084365822 | 0.000908681 |
| NM_001107763 | Ltk | leukocyte receptor tyrosine kinase | 4.018737034 | 0.023537516 |
| NM_021589 | Ntrk1 | neurotrophic receptor tyrosine kinase 1 | 3.855032183 | 0.024274736 |
| NM_001109419 | Apoc4 | apolipoprotein C4 | 3.813419178 | 0.005206018 |
| NM_001191822 | Rhbdl1 | rhomboid like 1 | 3.498096885 | 0.006871344 |
| NM_172047 | Eaf2 | ELL associated factor 2 | 3.470704647 | 0.013662039 |
| NM_001134562 | Rasal3 | RAS protein activator like 3 | 3.462527467 | 0.01411219 |
| NM_001107572 | Ovol1 | ovo like transcriptional repressor 1 | 3.372839189 | 0.013790267 |
| NM_001024360 | Lrrc73 | | 3.294792389 | 0.011273135 |
| NM_022234 | Asic4 | acid sensing ion channel subunit family member 4 | 3.289974713 | 0.048996677 |
| NM_017193 | Aadat | aminoadipate aminotransferase | 3.254592587 | 0.032884732 |
| NM_001002804 | C1rl | complement C1r subcomponent like | 3.198154658 | 0.031540286 |
| NM_001301812,N | Crhr1 | corticotropin releasing hormone receptor 1 | 3.173932237 | 0.02947819 |
| NM_031525 | Pdgfrb | platelet derived growth factor receptor beta | 3.155538577 | 9.50E-06 |
| NM_001109049 | Ccdc103 | coiled-coil domain containing 103 | 3.136847761 | 0.044093415 |
| NM_133298 | Gpnmb | glycoprotein nmb | 3.10844331 | 7.89E-05 |
| NM_001127540 | Lmntd2 | lamin tail domain containing 2 | 3.050451149 | 0.003044703 |
| NM_017250 | Htr2b | 5-hydroxytryptamine receptor 2B | 2.983068279 | 0.015934826 |
| NM_031056 | Mmp14 | matrix metallopeptidase 14 | 2.950376309 | 0.000231049 |
| NM_001108469 | Cdc42ep5 | CDC42 effector protein 5 | 2.92404232 | 0.005596293 |
| NM_001025772 | Stpg1 | sperm-tail PG-rich repeat containing 1 | 2.918903105 | 0.007027428 |
| NM_001277264,N | Apoa5 | apolipoprotein A5 | 2.748737726 | 0.015127821 |
| NM_053356,NM_0 | Col1a2 | collagen type I alpha 2 chain | 2.748650951 | 0.030197987 |
| NM_030868 | Nov | cellular communication network factor 3 | 2.736157254 | 0.002250091 |
| NM_001024342 | Dnai1 | dynein, axonemal, intermediate chain 1 | 2.735957272 | 0.002344052 |
| NM_019370 | Enpp3 | ectonucleotide pyrophosphatase/phosphodiesterase 3 | 2.670984226 | 0.024219051 |
| NR_024118 | Tnxa-ps1 | | 2.661907108 | 0.031623118 |
| NM_001107623 | Tmem26 | transmembrane protein 26 | 2.650181119 | 0.015224534 |
| NM_001137622 | Adamts2 | ADAM metallopeptidase with thrombospondin type 1 motif, 2 | 2.649746367 | 0.03013333 |
| NM_001014043 | Sgms2 | sphingomyelin synthase 2 | 2.642897195 | 0.025468204 |
| NM_001017514 | Izumo1 | izumo sperm-egg fusion 1 | 2.58878294 | 0.004952221 |
| NM_001108816 | Piga | phosphatidylinositol glycan anchor biosynthesis, class A | 2.551822261 | 0.003771704 |
| NM_001106777 | RGD1561102 | similar to ribosomal protein S12 | 2.527781034 | 0.039001418 |
| NM_001014041 | Fam46c | terminal nucleotidyltransferase 5C | 2.509155365 | 0.013181522 |

| | | | | |
|---|---|---|---|---|
| NM_001135600 | Cyp4v3 | cytochrome P450, family 4, subfamily v, polypeptide 3 | 2.488351317 | 0.006319003 |
| NM_001013171 | Gulp1 | GULP PTB domain containing engulfment adaptor 1 | 2.480867472 | 0.030827376 |
| NM_001002805 | C4b | complement C4B (Chido blood group) | 2.477015965 | 0.004774573 |
| NM_001013102 | Theg | theg spermatid protein | 2.430575423 | 0.002380573 |
| NM_001127541,N | Cracr2b | calcium release activated channel regulator 2B | 2.430109739 | 0.04603197 |
| NM_001013137 | Cxcl14 | C-X-C motif chemokine ligand 14 | 2.417848843 | 0.000194173 |
| NM_001106702 | Angptl6 | angiopoietin-like 6 | 2.382867386 | 0.005397008 |
| NM_001107182 | Crb1 | crumbs cell polarity complex component 1 | 2.380761092 | 0.007339484 |
| NM_001011984 | Asb2 | ankyrin repeat and SOCS box-containing 2 | 2.34818169 | 0.007910021 |
| NM_053338 | Rrad | RRAD, Ras related glycolysis inhibitor and calcium channel regulator | 2.345985376 | 0.020271895 |
| NM_031091 | Rab3b | RAB3B, member RAS oncogene family | 2.321269667 | 0.00021119 |
| NM_133559 | Pcsk4 | proprotein convertase subtilisin/kexin type 4 | 2.316766164 | 0.003306001 |
| NM_001013160 | Trim69 | tripartite motif-containing 69 | 2.30811084 | 0.015223422 |
| NM_001271115 | Dnaaf3 | dynein, axonemal, assembly factor 3 | 2.276079767 | 0.017373821 |
| NM_053955 | Crym | crystallin, mu | 2.271364757 | 0.007842237 |
| NM_022675 | Fkbp1b | FKBP prolyl isomerase 1B | 2.249641496 | 0.03546005 |
| NM_001108569 | Gbp5 | guanylate binding protein 5 | 2.199890284 | 0.03821303 |
| NM_012851 | Hsd17b1 | hydroxysteroid (17-beta) dehydrogenase 1 | 2.198366294 | 0.034091885 |
| NM_053442 | Slc7a8 | solute carrier family 7 member 8 | 2.181775538 | 0.001543119 |
| NM_001134736 | Wdr63 | WD repeat domain 63 | 2.181497882 | 0.017739466 |
| NM_001106231 | Ppm1n | protein phosphatase, Mg2+/Mn2+ dependent 1N | 2.166303157 | 0.010587944 |
| NM_001025044 | Ccdc146 | coiled-coil domain containing 146 | 2.149838594 | 0.010747091 |
| NM_001014068 | Hpdl | 4-hydroxyphenylpyruvate dioxygenase-like | 2.119886636 | 0.034994583 |
| NM_001107246 | Mfsd7 | solute carrier family 49 member 3 | 2.111358152 | 0.015000457 |
| NM_013180 | Itgb4 | integrin subunit beta 4 | 2.09286365 | 0.007606486 |
| NM_001013243 | Slc30a3 | solute carrier family 30 member 3 | 2.054755316 | 0.030171361 |
| NM_001012059 | Mcoln3 | mucolipin 3 | 2.053841022 | 0.007810665 |
| NM_001038615 | Fndc1 | fibronectin type III domain containing 1 | 2.034485922 | 0.04126362 |
| NM_001008880 | Scn4b | sodium voltage-gated channel beta subunit 4 | 2.010282603 | 0.035700654 |
| NM_001100741 | Col6a2 | collagen type VI alpha 2 chain | 2.003400756 | 0.000663071 |
| NM_001134695 | Fam229b | | 2.002878271 | 0.001449445 |
| NM_175592 | Cacna2d2 | calcium voltage-gated channel auxiliary subunit alpha2delta 2 | 1.996318288 | 0.031750248 |
| NM_053870 | Kcnj4 | potassium inwardly-rectifying channel, subfamily J, member 4 | 1.991211741 | 0.036379663 |
| NM_001109226 | Prrt4 | proline-rich transmembrane protein 4 | 1.985564856 | 0.029506408 |
| NM_001271179 | Slc25a45 | solute carrier family 25, member 45 | 1.985034297 | 0.027917631 |
| NM_001105800 | Zmynd15 | zinc finger, MYND-type containing 15 | 1.973386679 | 0.041187248 |
| NM_001109585 | Trim47 | tripartite motif-containing 47 | 1.968640757 | 0.044714619 |
| NM_012912 | Atf3 | activating transcription factor 3 | 1.960600604 | 0.038414275 |
| NM_057201 | Gpr37 | G protein-coupled receptor 37 | 1.948128409 | 0.035046166 |
| NM_021680 | Nxph4 | neurexophilin 4 | 1.944355321 | 0.023273056 |
| NM_001010965 | Mok | MOK protein kinase | 1.938018071 | 0.034434393 |
| NM_134432 | Agt | angiotensinogen | 1.931336455 | 8.33E-05 |
| NM_001108226 | Wnt6 | Wnt family member 6 | 1.917570778 | 0.00254144 |
| NM_181380 | Rtn4rl2 | reticulon 4 receptor-like 2 | 1.909858343 | 0.006000854 |
| NM_053500 | Slc25a27 | solute carrier family 25, member 27 | 1.905523196 | 0.030927106 |
| NM_001106348 | Mlana | melan-A | 1.885160314 | 0.001502623 |
| NM_001108227 | Wnt10a | Wnt family member 10A | 1.884199995 | 0.030626645 |
| NM_019386 | Tgm2 | transglutaminase 2 | 1.877021487 | 0.043717816 |
| NM_001008724,N | Fga | fibrinogen alpha chain | 1.869408476 | 0.039353688 |
| NM_001172103 | Pim2 | | 1.863151801 | 0.005622438 |
| NM_001271272 | Ndufa4l2 | NDUFA4, mitochondrial complex associated like 2 | 1.862467872 | 0.034804663 |
| NM_001109655 | Car14 | carbonic anhydrase 14 | 1.85928131 | 0.027378896 |
| NM_001107767 | Duoxa1 | dual oxidase maturation factor 1 | 1.85857763 | 0.021093079 |
| NM_001107857 | Ephb6 | Eph receptor B6 | 1.850186624 | 0.010932276 |
| NM_012676 | Tnnt2 | troponin T2, cardiac type | 1.848115508 | 0.041574584 |
| NR_130129 | LOC104940696 | | 1.844136133 | 0.00428513 |
| NM_001172079 | Icam5 | intercellular adhesion molecule 5 | 1.841237579 | 0.004512566 |
| NM_001109227 | Tspan33 | tetraspanin 33 | 1.833629547 | 0.004641388 |
| NM_001014051 | Ttll9 | tubulin tyrosine ligase like 9 | 1.829841639 | 0.014392166 |
| NM_001013072 | Sfxn2 | sideroflexin 2 | 1.817197797 | 0.018140056 |
| NM_001077680 | Bpifb1 | BPI fold containing family B, member 1 | 1.812205628 | 0.006398882 |
| NM_199085 | Serpinb6 | serpin family B member 6A | 1.805367245 | 0.021569098 |
| NM_013166 | Cntf | ciliary neurotrophic factor | 1.792471147 | 0.008812104 |
| NM_031504 | C4a | complement C4A | 1.784968478 | 0.000609862 |
| NM_001001514,N | Ablim2 | actin binding LIM protein family, member 2 | 1.775598327 | 0.004350853 |
| NM_019363 | Aox1 | aldehyde oxidase 1 | 1.762008497 | 0.001723391 |
| NM_001271346 | Ddb2 | damage specific DNA binding protein 2 | 1.745584361 | 0.009380517 |
| NM_001308302,N | Cacna1g | calcium voltage-gated channel subunit alpha1 G | 1.744346994 | 0.000546722 |
| NM_001128152 | Rwdd3 | RWD domain containing 3 | 1.743366983 | 0.014968507 |
| NM_001109574 | Tmem169 | transmembrane protein 169 | 1.740324097 | 0.007131359 |
| NM_013015 | Ptgds | prostaglandin D2 synthase | 1.731518979 | 0.048804518 |
| NM_080688 | Plcd4 | phospholipase C, delta 4 | 1.728447396 | 0.018280326 |
| NM_001270681,N | Apoe | apolipoprotein E | 1.721278189 | 0.000340202 |
| NM_001107724 | Tram1l1 | translocation associated membrane protein 1-like 1 | 1.719045671 | 0.026881683 |
| NM_012868 | Npr3 | natriuretic peptide receptor 3 | 1.71065961 | 0.001402506 |
| NM_031548 | Scnn1a | sodium channel epithelial 1 subunit alpha | 1.707042679 | 0.03063938 |
| NM_001270855,N | Stmn4 | stathmin 4 | 1.701357133 | 0.028656736 |
| NM_001134845 | LOC688613 | hypothetical protein LOC688613 | 1.695775326 | 0.01551456 |

| | | | | |
|---|---|---|---|---|
| NM_001276304,N | Dlgap3 | DLG associated protein 3 | 1.69455377 | 0.009316655 |
| NM_181636 | Col23a1 | collagen type XXIII alpha 1 chain | 1.690521862 | 0.019923778 |
| NM_031597 | Kcnq3 | potassium voltage-gated channel subfamily Q member 3 | 1.683511045 | 0.035645631 |
| NM_012650 | Shbg | sex hormone binding globulin | 1.677562282 | 0.043817765 |
| NM_001108978 | Pik3cd | phosphatidylinositol-4,5-bisphosphate 3-kinase, catalytic subunit delta | 1.671466662 | 0.032534986 |
| NM_001109160 | Flrt1 | fibronectin leucine rich transmembrane protein 1 | 1.664846035 | 0.012806989 |
| NM_001106713 | Klhl29 | | 1.662999796 | 0.023035955 |
| NM_001106877 | Rfx2 | regulatory factor X2 | 1.661784625 | 0.019386213 |
| NM_001107248 | Vcl | vinculin | 1.64233194 | 0.003558683 |
| NM_017014 | Gstm1 | glutathione S-transferase mu 1 | 1.641508853 | 0.006104227 |
| NM_001025048 | Setmar | SET domain and mariner transposase fusion gene | 1.640641275 | 0.04200108 |
| NM_001009709 | Tmem140 | transmembrane protein 140 | 1.635276991 | 0.023632814 |
| NM_053304 | Col1a1 | collagen type I alpha 1 chain | 1.619038929 | 0.024643617 |
| NM_001191778 | Aldh1l2 | aldehyde dehydrogenase 1 family, member L2 | 1.611367111 | 0.029039592 |
| NM_130411 | Coro1a | coronin 1A | 1.607898287 | 0.027062065 |
| NM_031154 | Gstm7 | glutathione S-transferase, mu 7 | 1.590868524 | 0.003336554 |
| NM_001108065 | Shc2 | SHC adaptor protein 2 | 1.577211924 | 0.041894813 |
| NM_001160162,N | Scn5a | sodium voltage-gated channel alpha subunit 5 | 1.574973314 | 0.011586839 |
| NM_019161 | Cdh22 | cadherin 22 | 1.568801176 | 0.025028534 |
| NM_001106276 | Cpeb1 | cytoplasmic polyadenylation element binding protein 1 | 1.565659685 | 0.002065137 |
| NM_133606 | Ehhadh | enoyl-CoA hydratase and 3-hydroxyacyl CoA dehydrogenase | 1.564130814 | 0.03642322 |
| NM_023981 | Csf1 | colony stimulating factor 1 | 1.56389721 | 0.002249108 |
| NM_001011976 | Wdr31 | | 1.558293089 | 0.027430515 |
| NM_017009 | Gfap | glial fibrillary acidic protein | 1.550004168 | 0.004017337 |
| NM_001030042 | Rad9b | RAD9 checkpoint clamp component B | 1.541249498 | 0.019168947 |
| NM_001109183 | Lhfp | LHFPL tetraspan subfamily member 6 | 1.533454143 | 0.002255867 |
| NM_138858 | Slc9a5 | solute carrier family 9 member A5 | 1.524382113 | 0.033650301 |
| NM_001108098 | Cpm | carboxypeptidase M | 1.517368103 | 0.008216459 |
| NM_001017478 | Cxcl16 | C-X-C motif chemokine ligand 16 | 1.51671187 | 0.012903395 |
| NM_080782 | Cdkn1a | cyclin-dependent kinase inhibitor 1A | 1.512067924 | 0.029627177 |
| NM_001130499 | Ttc38 | | 1.510134741 | 0.036211686 |
| NM_001108987 | Diras1 | DIRAS family GTPase 1 | 1.509655659 | 0.042750169 |

**Gene list whose expression level is <0.8-fold reduced in siDerl1 NSCs**

| Refseq ID | GeneSymbol | Description | Fold change | P value |
|---|---|---|---|---|
| NM_001017510 | LOC498750 | | 0.011912108 | 0.007776697 |
| NM_031560 | Ctsk | cathepsin K | 0.022471789 | 0.000146434 |
| NM_012841 | Dcc | DCC netrin 1 receptor | 0.026872545 | 0.002358742 |
| NM_001107671 | Plcxd3 | phosphatidylinositol-specific phospholipase C, X domain containing 3 | 0.032147715 | 0.002587457 |
| NM_001108996 | Ap1m2 | adaptor related protein complex 1 subunit mu 2 | 0.033645063 | 0.014430631 |
| NM_001024907 | MGC114499 | | 0.044566363 | 0.029718305 |
| NM_001191843 | Atp6ap1l | ATPase H+ transporting accessory protein 1 like | 0.046196993 | 0.012600929 |
| NM_001105737 | Tek | TEK receptor tyrosine kinase | 0.047187309 | 0.01270479 |
| NM_133420 | Chrna2 | cholinergic receptor nicotinic alpha 2 subunit | 0.051363696 | 0.04169532 |
| NM_022636 | Vax1 | ventral anterior homeobox 1 | 0.052576863 | 0.026075399 |
| NM_138897 | Gabrr3 | gamma-aminobutyric acid type A receptor rho3 subunit | 0.053784985 | 0.026759072 |
| NM_001191728 | Lancl3 | LanC like 3 | 0.054034641 | 0.046316639 |
| NM_053423 | Tert | telomerase reverse transcriptase | 0.060818771 | 0.046825904 |
| NM_001109130 | Tsnaxip1 | translin-associated factor X interacting protein 1 | 0.061169113 | 0.039675912 |
| NM_053934 | Pcdha13 | protocadherin alpha 13 | 0.061314119 | 0.047254662 |
| NM_001107056 | Cyb561 | cytochrome b-561 | 0.061762371 | 0.04386896 |
| NM_001109641 | Hist3h2bb | H2B.U histone 1 | 0.062856005 | 0.044362396 |
| NM_031796 | Galnt5 | polypeptide N-acetylgalactosaminyltransferase 5 | 0.072997826 | 0.00677792 |
| NM_031828 | Kcnma1 | potassium calcium-activated channel subfamily M alpha 1 | 0.094163072 | 0.000210535 |
| NM_053744 | Dlk1 | delta like non-canonical Notch ligand 1 | 0.103349815 | 0.003838884 |
| NM_001108051 | Slc24a4 | solute carrier family 24 member 4 | 0.109592958 | 0.000174471 |
| NM_012636 | Pthlh | parathyroid hormone-like hormone | 0.116185855 | 0.000506216 |
| NM_001100523 | Otp | orthopedia homeobox | 0.12013391 | 0.007099 |
| NM_001111114,N | Grik1 | glutamate ionotropic receptor kainate type subunit 1 | 0.120892009 | 0.031771457 |
| NM_001107755 | Pamr1 | peptidase domain containing associated with muscle regeneration 1 | 0.128402805 | 0.011777393 |
| NM_013120 | Gckr | glucokinase regulator | 0.129700017 | 0.01277519 |
| NM_022407 | Aldh1a1 | aldehyde dehydrogenase 1 family, member A1 | 0.139129865 | 0.001992054 |
| NM_001305138,N | Synpo2l | synaptopodin 2-like | 0.144837167 | 0.041216021 |
| NM_053318 | Hpx | hemopexin | 0.159402072 | 0.045434139 |
| NM_031741 | Slc2a5 | solute carrier family 2 member 5 | 0.17238264 | 0.03198011 |
| NM_023100 | Nmur1 | neuromedin U receptor 1 | 0.177634398 | 0.011291368 |
| NM_019190 | Cd46 | CD46 molecule | 0.185976367 | 0.027572512 |
| NM_031012 | Anpep | alanyl aminopeptidase, membrane | 0.207542493 | 0.021753365 |
| NM_138530 | Pbld1 | phenazine biosynthesis-like protein domain containing 1 | 0.22379207 | 0.004746682 |
| NM_001127650 | Anks4b | ankyrin repeat and sterile alpha motif domain containing 4B | 0.225229537 | 0.043559439 |
| NM_053977 | Cdh17 | cadherin 17 | 0.230005421 | 0.024744727 |
| NM_198748 | Scin | scinderin | 0.240124233 | 0.028676846 |
| NM_001108568 | Dapp1 | dual adaptor of phosphotyrosine and 3-phosphoinositides 1 | 0.240432876 | 0.003788887 |
| NM_031117 | Snrpn | small nuclear ribonucleoprotein polypeptide N | 0.250782252 | 0.031408865 |
| NM_001130502 | Fam83f | | 0.251396067 | 0.003843109 |

| | | | | |
|---|---|---|---|---|
| NM_031549 | Tagln | transgelin | 0.268809702 | 0.003157927 |
| NM_001033687 | Ushbp1 | USH1 protein network component harmonin binding protein 1 | 0.27018035 | 0.015159021 |
| NM_053441 | Slco1c1 | solute carrier organic anion transporter family, member 1c1 | 0.270762242 | 0.042160854 |
| NM_001109309 | Cdk5r2 | cyclin-dependent kinase 5 regulatory subunit 2 | 0.273806725 | 0.013966022 |
| NM_001106267 | Tm2d3 | TM2 domain containing 3 | 0.288605669 | 0.015286944 |
| NM_001077677 | Pacrg | parkin coregulated | 0.30723147 | 0.000451762 |
| NM_001127557 | Rtdr1 | | 0.314455776 | 0.010457361 |
| NM_032070 | Hmga2 | high mobility group AT-hook 2 | 0.334480841 | 0.000995459 |
| NM_001076553,N | Pkib | cAMP-dependent protein kinase inhibitor beta | 0.350935159 | 0.001112807 |
| NM_001008838 | RT1-CE15 | | 0.36549479 | 0.031009856 |
| NM_001047914 | Sdhaf3 | succinate dehydrogenase complex assembly factor 3 | 0.372738155 | 0.01523939 |
| NM_001191694 | Nebl | nebulette | 0.373426558 | 0.008008049 |
| NM_031347 | Ppargc1a | PPARG coactivator 1 alpha | 0.377660729 | 0.01916138 |
| NM_133285 | Hist1h1d | H1.4 linker histone, cluster member | 0.382868558 | 0.008112707 |
| NM_001127602 | Slc25a53 | | 0.383742202 | 0.000379422 |
| NM_001108859 | Irf6 | interferon regulatory factor 6 | 0.400224974 | 0.00349448 |
| NM_019265 | Scn11a | sodium voltage-gated channel alpha subunit 11 | 0.400917748 | 0.005851973 |
| NM_001009695 | Wnt7b | Wnt family member 7B | 0.403075487 | 0.031225081 |
| NM_001127640 | PCOLCE2 | procollagen C-endopeptidase enhancer 2 | 0.406032106 | 0.000170092 |
| NM_138524 | A3galt2 | alpha 1,3-galactosyltransferase 2 | 0.418470472 | 0.000934804 |
| NM_001024282 | Hist1h2af | histone cluster 1 H2A family member F | 0.42040864 | 0.044597569 |
| NM_173136 | Akr1b8 | aldo-keto reductase family 1, member B8 | 0.421725801 | 0.005579554 |
| NM_001012460 | Septin1 | septin 1 | 0.422672974 | 0.030943282 |
| NM_001107170 | Tfcp2l1 | transcription factor CP2-like 1 | 0.431626543 | 0.012843802 |
| NM_021658 | Hcn4 | hyperpolarization activated cyclic nucleotide-gated potassium channel 4 | 0.433367586 | 0.022050108 |
| NM_024141 | Duox2 | dual oxidase 2 | 0.443565483 | 0.002512099 |
| NM_053294 | Adora2a | adenosine A2a receptor | 0.448932684 | 0.006610831 |
| NM_001135710 | Sec14l5 | | 0.45319148 | 0.023124069 |
| NM_053544 | Sfrp4 | secreted frizzled-related protein 4 | 0.464254873 | 0.004762944 |
| NM_080894 | Pde7b | phosphodiesterase 7B | 0.466425359 | 0.002076483 |
| NM_001107400 | Celf4 | CUGBP, Elav-like family member 4 | 0.471613245 | 0.019419984 |
| NM_053882 | Ecm1 | extracellular matrix protein 1 | 0.473367502 | 2.55E-07 |
| NM_001033961,N | Kcnip2 | potassium voltage-gated channel interacting protein 2 | 0.482784906 | 0.013140678 |
| NM_030875 | Scn1a | sodium voltage-gated channel alpha subunit 1 | 0.487866666 | 0.009273831 |
| NM_001107533 | Adamtsl3 | ADAMTS-like 3 | 0.491757199 | 0.04063176 |
| NM_175595 | Cacna2d3 | calcium voltage-gated channel auxiliary subunit alpha2delta 3 | 0.493075025 | 0.017503562 |
| NM_031720 | Dio2 | iodothyronine deiodinase 2 | 0.50477183 | 0.007950121 |
| NM_001105884 | Hoxd1 | homeo box D1 | 0.505886421 | 0.022356274 |
| NM_001108750 | Cpne8 | copine 8 | 0.508941304 | 0.013103961 |
| NM_019345 | Slc12a3 | solute carrier family 12 member 3 | 0.512209036 | 0.049055449 |
| NM_001190459 | LOC100361087 | hypothetical LOC100361087 | 0.518491737 | 0.039681662 |
| NM_001007726 | Dnai2 | dynein, axonemal, intermediate chain 2 | 0.523199014 | 0.009104589 |
| NM_001108061 | Amn | amnion associated transmembrane protein | 0.54810817 | 0.038741379 |
| NM_001276707,N | Srebf1 | sterol regulatory element binding transcription factor 1 | 0.556245434 | 0.011812783 |
| NM_001008320 | Rhoj | ras homolog family member J | 0.580693316 | 0.011184995 |
| NM_001108881 | Rnf144b | ring finger protein 144B | 0.585156181 | 0.011539822 |
| NM_199105 | Fam198b | golgi associated kinase 1B | 0.58923031 | 0.000228659 |
| NM_024371 | Slc6a1 | solute carrier family 6 member 1 | 0.589839295 | 0.009989473 |
| NM_001100512 | Gpld1 | glycosylphosphatidylinositol specific phospholipase D1 | 0.597979202 | 0.019668805 |
| NM_001112716,N | Grik3 | glutamate ionotropic receptor kainate type subunit 3 | 0.598879544 | 0.010845673 |
| NM_012994 | Nxph1 | neurexophilin 1 | 0.599853605 | 0.001405973 |
| NM_053352 | Ackr3 | atypical chemokine receptor 3 | 0.600064841 | 0.006864116 |
| NM_001012215 | Pcdhgb7 | protocadherin gamma subfamily B, 7 | 0.605833803 | 0.037492738 |
| NM_173151 | Pcyt1b | phosphate cytidylyltransferase 1, choline, beta | 0.618230805 | 0.032019948 |
| NM_001107949 | Dnajc6 | DnaJ heat shock protein family (Hsp40) member C6 | 0.620788341 | 0.029973309 |
| NM_031779 | Apba1 | amyloid beta precursor protein binding family A member 1 | 0.629392479 | 0.018761387 |
| NM_001108036 | Plekhh1 | pleckstrin homology, MyTH4 and FERM domain containing H1 | 0.63146503 | 0.025616984 |
| NM_031672 | Slc15a2 | solute carrier family 15 member 2 | 0.636243788 | 0.045034058 |
| NM_001108897 | Pskh1 | protein serine kinase H1 | 0.640305324 | 0.012230094 |
| NM_001109333 | Lrrc14b | leucine rich repeat containing 14B | 0.641081459 | 0.042831529 |
| NM_001191757 | Frem1 | Fras1 related extracellular matrix 1 | 0.645182518 | 0.008141812 |
| NM_199504 | Pcdha2 | protocadherin alpha 2 | 0.645954922 | 0.032206219 |
| NM_023025 | Cyp2j4 | cytochrome P450, family 2, subfamily j, polypeptide 4 | 0.669988788 | 0.015458869 |
| NM_001106578 | Sema3c | semaphorin 3C | 0.675334069 | 0.027766582 |
| NM_001270807,N | Sphk1 | sphingosine kinase 1 | 0.676104448 | 0.046936434 |
| NM_001105932 | Vps29 | VPS29 retromer complex component | 0.676354549 | 0.000360785 |
| NM_001114602 | Pcdhb5 | protocadherin beta 5 | 0.679670139 | 0.012462884 |
| NM_001004277 | Pla2g15 | phospholipase A2, group XV | 0.684140728 | 0.013277384 |
| NM_001106648 | Rpp25l | ribonuclease P/MRP subunit p25 like | 0.684741275 | 0.009506729 |
| NM_001008292 | Diablo | diablo, IAP-binding mitochondrial protein | 0.685857547 | 0.000179142 |
| NM_053986 | Myo1b | myosin Ib | 0.68883597 | 0.041102114 |
| NM_001109278 | Isca2 | iron-sulfur cluster assembly 2 | 0.692051787 | 0.035691068 |
| NM_001106668 | Slc35d1 | solute carrier family 35 member D1 | 0.694338971 | 0.019767079 |
| NM_001007802 | Cmtm6 | CKLF-like MARVEL transmembrane domain containing 6 | 0.701815684 | 0.038073015 |
| NM_173094 | Hmgcs2 | 3-hydroxy-3-methylglutaryl-CoA synthase 2 | 0.704058672 | 0.033591612 |
| NM_001191821 | Frmd4a | FERM domain containing 4A | 0.705340341 | 0.004320738 |
| NM_001289778 | Map7d2 | MAP7 domain containing 2 | 0.705621497 | 0.035780989 |
| NM_001271026 | Zcchc10 | zinc finger CCHC-type containing 10 | 0.71239681 | 0.034268636 |

| | | | | |
|---|---|---|---|---|
| NM_017218 | Erbb3 | erb-b2 receptor tyrosine kinase 3 | 0.712891704 | 0.009875152 |
| NM_023989 | Senp2 | SUMO specific peptidase 2 | 0.713996586 | 0.013879892 |
| NM_001106095 | Lig4 | DNA ligase 4 | 0.714377204 | 0.019108192 |
| NM_030994 | Itga1 | integrin subunit alpha 1 | 0.714391939 | 0.036724595 |
| NM_024154 | Asic1 | acid sensing ion channel subunit 1 | 0.717460566 | 0.014014056 |
| NM_053725 | Itga6 | integrin subunit alpha 6 | 0.718220959 | 0.000573662 |
| NM_001109014 | Tmem164 | transmembrane protein 164 | 0.721948612 | 0.034035998 |
| NM_013119 | Scn3a | sodium voltage-gated channel alpha subunit 3 | 0.722367879 | 0.021964174 |
| NM_001107275 | Slc39a14 | solute carrier family 39 member 14 | 0.724295488 | 0.024309902 |
| NM_053886 | Lman1 | lectin, mannose-binding, 1 | 0.727318009 | 0.010286139 |
| NM_053379 | Dcx | doublecortin | 0.729502957 | 0.006864881 |
| NM_012802 | Pdgfra | platelet derived growth factor receptor alpha | 0.730678335 | 0.038514844 |
| NM_017206 | Slc6a6 | solute carrier family 6 member 6 | 0.731732421 | 0.004388351 |
| NM_001107596 | Tmem2 | cell migration inducing hyaluronidase 2 | 0.734671323 | 0.033125403 |
| NM_019147 | Jag1 | jagged canonical Notch ligand 1 | 0.741250877 | 0.008053911 |
| NM_201990 | Pgap1 | post-GPI attachment to proteins inositol deacylase 1 | 0.742931636 | 0.038386969 |
| NM_001014076 | Nol10 | nucleolar protein 10 | 0.743576006 | 0.011393955 |
| NM_001012201 | Cadm1 | cell adhesion molecule 1 | 0.745122006 | 0.002454913 |
| NM_001127527 | Yeats4 | YEATS domain containing 4 | 0.746002878 | 0.010673263 |
| NM_022380 | Stat5b | signal transducer and activator of transcription 5B | 0.751412835 | 0.009365889 |
| NM_001271344 | Lpcat2 | lysophosphatidylcholine acyltransferase 2 | 0.751488425 | 0.039880658 |
| NM_017022 | Itgb1 | integrin subunit beta 1 | 0.751658059 | 0.000656344 |
| NM_053943 | Pcdhgc3 | protocadherin gamma subfamily C, 3 | 0.752165614 | 0.001079378 |
| NM_001100863 | Galnt16 | polypeptide N-acetylgalactosaminyltransferase 16 | 0.752214747 | 0.02731447 |
| NM_001108892 | Sall3 | spalt-like transcription factor 3 | 0.753699165 | 0.025606453 |
| NM_001107548 | Usp31 | ubiquitin specific peptidase 31 | 0.755223977 | 0.030982569 |
| NM_001013990 | Srd5a3 | steroid 5 alpha-reductase 3 | 0.760789945 | 0.015359354 |
| NM_024486 | Acvr1 | activin A receptor type 1 | 0.761452399 | 0.028917579 |
| NM_001126283 | Ankrd49 | ankyrin repeat domain 49 | 0.763885647 | 0.027363235 |
| NM_053467 | Tmed10 | transmembrane p24 trafficking protein 10 | 0.765917187 | 0.002965834 |
| NM_199407 | Unc5c | unc-5 netrin receptor C | 0.766019514 | 0.035271861 |
| NM_001006980 | Bcap29 | B-cell receptor-associated protein 29 | 0.767537301 | 0.027649899 |
| NM_001128079 | Ctdsp1 | CTD small phosphatase 1 | 0.768993761 | 0.004059327 |
| NM_001107374 | RGD1308601 | similar to hypothetical protein | 0.76949287 | 0.031424918 |
| NM_031800 | Dedd | death effector domain-containing | 0.770677262 | 0.047690615 |
| NM_013057 | F3 | coagulation factor III, tissue factor | 0.771051135 | 0.027718463 |
| NM_001173972 | Itga8 | integrin subunit alpha 8 | 0.771638824 | 0.022346724 |
| NM_001015027 | Crebl2 | cAMP responsive element binding protein-like 2 | 0.771934622 | 0.024998338 |
| NM_001171177 | Tmtc2 | transmembrane O-mannosyltransferase targeting cadherins 2 | 0.772256114 | 0.009702114 |
| NM_001191704 | Fndc3b | fibronectin type III domain containing 3B | 0.775460136 | 0.018101049 |
| NM_001173472 | RGD1562987 | | 0.778326151 | 0.022140786 |
| NM_001025118 | Fam63a | MINDY lysine 48 deubiquitinase 1 | 0.778336803 | 0.047343985 |
| NM_053883 | Dusp6 | dual specificity phosphatase 6 | 0.779737615 | 0.039424181 |
| NM_001037139 | Pcdhga2 | protocadherin gamma subfamily A, 2 | 0.782777366 | 0.019784649 |
| NM_001111127 | Hist3h2ba | histone cluster 3, H2ba | 0.783624983 | 0.025444052 |
| NM_012880 | Sod3 | superoxide dismutase 3 | 0.78420287 | 0.030056506 |
| NM_080904_1 | Arf3 | | 0.784775936 | 0.006176481 |
| NM_001024303 | Lix1l | limb and CNS expressed 1 like | 0.785039391 | 0.006945989 |
| NM_022624 | Slc22a23 | solute carrier family 22, member 23 | 0.787541463 | 0.026849625 |
| NM_001108390 | Ndfip2 | Nedd4 family interacting protein 2 | 0.788380182 | 0.022321693 |
| NM_053502 | Abcg1 | ATP binding cassette subfamily G member 1 | 0.788567648 | 0.042874501 |
| NM_053948 | Polr2g | RNA polymerase II subunit G | 0.788991936 | 0.033518327 |
| NM_001191715 | Slc30a7 | solute carrier family 30 member 7 | 0.790483615 | 0.03924914 |
| NM_001005907,N | Efemp2 | EGF containing fibulin extracellular matrix protein 2 | 0.790985604 | 0.028163392 |
| NM_001037979 | Adipor2 | adiponectin receptor 2 | 0.79209672 | 0.041523386 |
| NM_001127639 | Gba | glucosylceramidase beta | 0.793632646 | 0.032446844 |
| NM_001037159 | Pcdhgb8 | protocadherin gamma subfamily B, 8 | 0.794921326 | 0.020106543 |
| NM_001007641 | Rnd3 | Rho family GTPase 3 | 0.79509397 | 0.041960276 |
| NR_130147,NR_13 | Tug1 | | 0.795641052 | 0.011961472 |
| NM_001108747 | Rassf3 | Ras association domain family member 3 | 0.797306936 | 0.044655465 |
| NM_134356 | Ptprg | protein tyrosine phosphatase, receptor type, G | 0.799474486 | 0.029473699 |
| NM_021266 | Fzd1 | frizzled class receptor 1 | 0.799715791 | 0.015379829 |

### Supplemental Materials and Methods

*DNA microarray analysis*

Total RNA was extracted from the DG using a NucleoSpin RNA kit (740955, Takara Bio) following the manufacturer's instructions. A total of 150 ng of total RNA from each sample was amplified and labeled with Cy3. Next, 600 ng Cy3-labeled cRNA was fragmented, hybridized onto the SurePrint G3 Mouse GE Ver2 platform (G4852B, Agilent Technologies), and then incubated at 65°C while being rotated for 17 h. Data were analyzed using GeneSpring software version 14.9 (Agilent Technologies) as previously described (Komatsu et al. 2013). In brief, the microarray data were normalized by quantile normalization, and the baseline signal values were transformed to the median in all samples. Then, quality control and filtering steps were performed based on flags and expression levels. Mean signal intensities were measured in duplicate and averaged to identify genes differentially expressed among mouse lines. Data from this microarray analysis have been submitted to the NCBI Gene Expression Omnibus archive as series GSE229342. GSEA was performed using GSEA v4.1.0 (https://www.gsea-msigdb.org/gsea/index.jsp). The enrichment plot shows the distribution of genes in each set that are positively (red) and negatively (blue) correlated with Derlin-1 deficiency. The Gene Ontology (GO) terms for GSEA were obtained from the Mouse Genome Informatics (MGI) GO project (http://www.informatics.jax.org/), which provides functional annotations for mouse gene products using Gene Ontology (http://www.informatics.jax.org/vocab/gene_ontology).

*RNA-seq*

RNA library construction and RNA-seq were performed using an Illumina sequencing platform (GENEWIZ). *Derl1* or its control knockdown adult rat-derived hippocampal NSCs were cultured in the presence of BMP4 for 2 days. Then, three samples of total RNA from the two cell groups were extracted for transcriptome sequencing and RNA-seq analysis. The cDNA libraries were used to construct the transcriptome sequence library in GENEWIZ (S. Plainfield, NJ) company using Illumina HiSeq X. The files containing the results were processed with a standard pipeline that included end trimming with trimommatic (Bolger et al. 2014). Then, the sequence reads were mapped to the rat reference genome (rn6) using STAR (Dobin et al. 2013). The mapped sequences were converted to expression levels (transcripts per million, TPM) and quantified using RSEM (Li and Dewey 2011). Differential gene expression analysis was performed using edgeR (Robinson et al. 2010). A fold change <0.8 was considered downregulation, and a fold change >1.5 was considered upregulation. Data from this RNA-seq analysis have been submitted to the NCBI Gene Expression Omnibus archive as series GSE229251. GSEA was performed using GSEA v4.1.0. The enrichment plot shows the distribution of genes in each set that are positively (red) and negatively (blue) correlated with *Derl1* knockdown. The GO terms for GSEA were obtained from the Rat Genome Database (RGD) (https://rgd.mcw.edu/), which provides functional annotations for rat gene products using Gene Ontology (https://rgd.mcw.edu/GO/). Stat5b target genes were obtained by searching ChIP-Atlas (https://chip-atlas.org), and the binding criterion was set to a maximum distance of 1 kb in either direction from the transcription start site of the target. After ChIP-Atlas scoring, the top 80 potential targets were used as the gene set for GSEA.

**Supplemental Table S2. List of primary and secondary antibodies used in this study.**

| Antibodies | SOURCE | IDENTIFIER |
|---|---|---|
| Mouse-anti-Derlin-1 | Sigma-Aldrich | SAB4200148; RRID: AB_10624068 |
| Rat-anti-BrdU | AbD Serotec | OBT0030; RRID: AB_609568 |
| Goat-anti-DCX | Santa Cruz | sc-8066; RRID: AB_2088494 |
| Rabbit-anti-Prox1 | Millipore | AB5475; RRID: AB_177485 |
| Mouse-anti-NeuN | Millipore | MAB377; RRID: AB_2298772 |
| Mouse-anti-Ki67 | BD PharMingen | 550609; RRID: AB_393778 |
| Rabbit-anti-Tbr2 | Abcam | ab23345; RRID: AB_778267 |
| Mouse-anti-Nestin | Millipore | MAB353; RRID: AB_94911 |
| Rabbit-anti-MCM2 | Cell Signaling | 3619; RRID: AB_2142137 |
| Chicken-anti-GFAP | Millipore | AB5541; RRID: AB_177521 |
| Goat-anti-Sox2 | Santa Cruz | Sc-17320; RRID: AB_2286684 |
| Rabbit-anti-HA-Tag | Cell Signaling | 3724; RRID: AB_1549585 |
| Mouse-anti-Stat5b | Santa Cruz | Sc1656-; RRID: AB_2197067 |
| Mouse-anti-Actin | Sigma–Aldrich | A4700; RRID: AB_476730 |
| Anti-Mouse IgG, HRP-linked antibody | GE Healthcare | NA931; RRID: AB_772210 |
| CF®555, Donkey Anti-Mouse IgG (H+L), Highly Cross-Adsorbed | Biotium | 20037; RRID: AB_10559035 |
| CF®555, Donkey Anti-Rabbit IgG (H+L), Highly Cross-Adsorbed | Biotium | 20038; RRID: AB_10558011 |
| CF®488A, Donkey Anti-Chicken IgY (H+L), Highly Cross-Adsorbed | Biotium | 20166; RRID: AB_10854387 |
| CF®488A, Donkey Anti-Mouse IgG (H+L), Highly Cross-Adsorbed | Biotium | 20014; RRID: AB_10561327 |
| CF®568, Donkey Anti-Rat IgG (H+L), Highly Cross-Adsorbed | Biotium | 20092; RRID: AB_10559037 |
| CF®647, Donkey Anti-Goat IgG (H+L), Highly Cross-Adsorbed | Biotium | 20048; RRID: AB_10853455 |
